# DrugGym: A testbed for the economics of autonomous drug discovery

**DOI:** 10.1101/2024.05.28.596296

**Authors:** Michael Retchin, Yuanqing Wang, Kenichiro Takaba, John D. Chodera

## Abstract

Drug discovery is stochastic. The effectiveness of candidate compounds in satisfying design objectives is unknown ahead of time, and the tools used for prioritization—predictive models and assays—are inaccurate and noisy. In a typical discovery campaign, thousands of compounds may be synthesized and tested before design objectives are achieved, with many others ideated but deprioritized. These challenges are well-documented, but assessing potential remedies has been difficult. We introduce **DrugGym**, a framework for modeling the stochastic process of drug discovery. Emulating biochemical assays with realistic surrogate models, we simulate the progression from weak hits to sub-micromolar leads with viable ADME. We use this testbed to examine how different ideation, scoring, and decision-making strategies impact statistical measures of utility, such as the probability of program success within predefined budgets and the expected costs to achieve target candidate profile (TCP) goals. We also assess the influence of affinity model inaccuracy, chemical creativity, batch size, and multi-step reasoning. Our findings suggest that reducing affinity model inaccuracy from 2 to 0.5 pIC50 units improves budget-constrained success rates tenfold. DrugGym represents a realistic testbed for machine learning methods applied to the hit-to-lead phase. Source code is available at www.drug-gym.org.

## Introduction

Small molecule drug discovery is slow, costly, and prone to failure. Inefficiencies and bottlenecks plague every stage of discovery programs, as half of them fail to produce a candidate for preclinical development [1]. Even this achievement can take more than 4.5 years and $13.5M on average [1]. Should a candidate be declared, clinical success is unlikely [2]. Amortizing failures, the cost of a novel drug approval is estimated to be $2.9 billion [3]. These challenges motivate the need for efficient discovery of multiple development candidates that satisfy the target candidate profile (TCP) objectives rather than a single choice.

### Drug discovery is a stochastic process

Candidates for preclinical development are difficult to find because drug discovery programs are realizations of a stochastic process aiming to solve complex, multi-objective design criteria. Each discovery campaign attempts to meet these criteria through iterations of design, synthesis, and analysis. By the time the program is terminated, it may succeed or fail in doing so to various degrees. Since the outcomes of experiments performed on novel compounds are unknown *a priori*, drug hunters use predictive models and surrogate assays to prioritize compounds. Yet predictive models are inaccurate and assays are prone to measurement noise. The ability to realistically simulate the underlying stochastic process would enable a new science of drug campaign interventions. We could then ask: how do different strategies or models impact the probability of successfully achieving program goals given time and budget constraints? Conversely, how do those choices reduce the expected time and cost required to reach the goals? One could also pose tactical queries, such as which chemical space is most enriched for leads with progressable ADME properties, or how much time and treasure is worthwhile to be spent improving predictive models.

### Interventions on the econometrics of drug discovery are urgently needed yet intractable

Strategic decisions in discovery campaigns vary by sponsoring organization, therapeutic area, regulatory considerations, and expected market dynamics, influencing how costs and timelines are prioritized. An academic center may happily pursue a long-running discovery program if it came with a reduced budget but a high probability of success. For a sufficiently large market, a pharma company may accept costly experiments that minimize time to market [3]. A government-funded program to identify antivirals amid a pandemic might elect to accelerate at virtually unlimited cost [4]. There are enormous incentives to consider these tradeoffs econometrically, as the global prescription drug market will be $1.6 trillion annually by 2026 [5].

Pharma industry econometrics has been extensively scrutinized, finding steep declines in R&D efficiency [6, 7]. These are observational studies, so while they clearly demonstrate the problem, they are unable to substantiate solutions. Doing so would be too costly: a real-world test that brings one alternative decision-making scheme to completion (reaching program goal) would cost as much as doing things the usual way, with dubious payoff. That is just for one trial. Appropriately powering such an experiment in a randomized setting is impracticable.

Computational methods have been cited as a potential panacea for inefficiency, since they have a history of exponential efficiency gains [8, 9]. Applications include molecular design [10–17], retrosynthesis [18–21], virtual screening [22–30], and ADMET prediction [31, 32]. Many of these systems are already in use within real drug discovery programs [33–35]. Alongside these methods, significant effort has been organized around developing robust benchmarks for predictive models [36, 37]. Yet these benchmarks are generally designed to demonstrate faithfulness to a single metric, which does not directly address the true impact or utility for discovery programs. Even targeting a composite metric such as QED druglikeness [38], we are no closer to answering the strategic questions posed above, nor to efficiently searching the vast design space of decision heuristics [39].

### Useful simulations of discovery campaigns must be grounded in chemistry

Previous authors have proposed models of discovery campaigns to study interventions on R&D productivity [40–42]. Inferences range from estimates of employee productivity [40] to the feasibility of program underwriting strategies [42], backed by the statistics of hand-designed Markov processes. Yet these simulations are not grounded in chemistry. Instead, transition probabilities between program stages (like hit-finding and preclinical development) are determined by literature or reasonable speculation. This means they lack adequate granularity to assess ideation strategies, synthetic capabilities, predictive models, or decision-making procedures. Moreover, they lack realism. Actual programs are stymied by optimization plateaus [43, 44], activity cliffs [45], and multiple simultaneous objectives that may be anti-correlated [46, 47].

One straightforward way to model this process qualitatively is with a self-avoiding walk, proposed by John Delaney [48]. He observed that discovery programs avoid ideating the same molecule twice, and modeling programs with this simple property is sufficient to make meaningful comments on impacts of chemist resources and cost expectations, stopping rules, structure-activity relationship (SAR), and the difficulty of design objectives. In the two decades since publication, surprisingly little work has followed up on these ideas.

### Drug discovery campaigns involve a series of decisions aiming to achieve specified goals

At the outset of a small molecule drug discovery campaign, researchers outline their objectives for the program in a **target candidate profile (TCP)**: the desired properties of a lead candidate that could then progress into costly preclinical development studies (**Table 1**). The TCP objectives span affinity against the targeted protein (here, measured in pIC50 units) and other properties that relate how the molecule is absorbed, distributed, metabolized, and excreted (ADME) relevant to the pharmacokinetics of the molecule, such as solubility (commonly expressed as log S) and lipophilicity (log P). They may include functional properties, like penetrance of the blood-brain barrier, rate of renal clearance, or target class selectivity [49]. Researchers set upper and lower bounds that determine the ideal ranges of these properties [50]. They then identify molecular hits—chemical matter with weak affinity and poor ADME properties and which, nevertheless, defines the starting point for prosecuting a given discovery program. From here, researchers aim to improve each compound’s properties by making a succession of designs that introduce small changes into the chemical structures of promising molecules. But only choices that can be practically synthesized are feasible, a function of the available chemical building blocks and reaction repertoire [51].

**Table 1.** Target candidate profile (TCP) objectives used in this work. Associated methods used here for computing corresponding Oracles in DrugGym are noted.

| Objective | Units | Oracle Method | Dynamic Range | Thresholds |  |
| --- | --- | --- | --- | --- | --- |
|  |  |  |  | Acceptable | Ideal |
| Affinity | pIC50 | Docking [56] | [3, 11] [57] | [8, ∞) | [9, ∞) |
| Lipophilicity | Log P | RDKit [58] | [-1, 8] [59] | [0, 4] | [0, 3] |
| Solubility | Log S | Gradient Boosted Tree [60] | [-8, 1] [61] | [-4, 0] | [-3, 0] |

A typical discovery program by a human team will first concentrate on optimizing affinity, even if it means compromising other TCP objectives. When sufficient affinity is achieved, attention is diverted to optimizing the other properties [52]. Progress is uncertain, and programs may occasionally backtrack. Eventually, a molecule is found that satisfies all objectives, provided the program budget (limited in money, time, or both) is not exhausted (**Figure 1A**).

**Figure 1.**
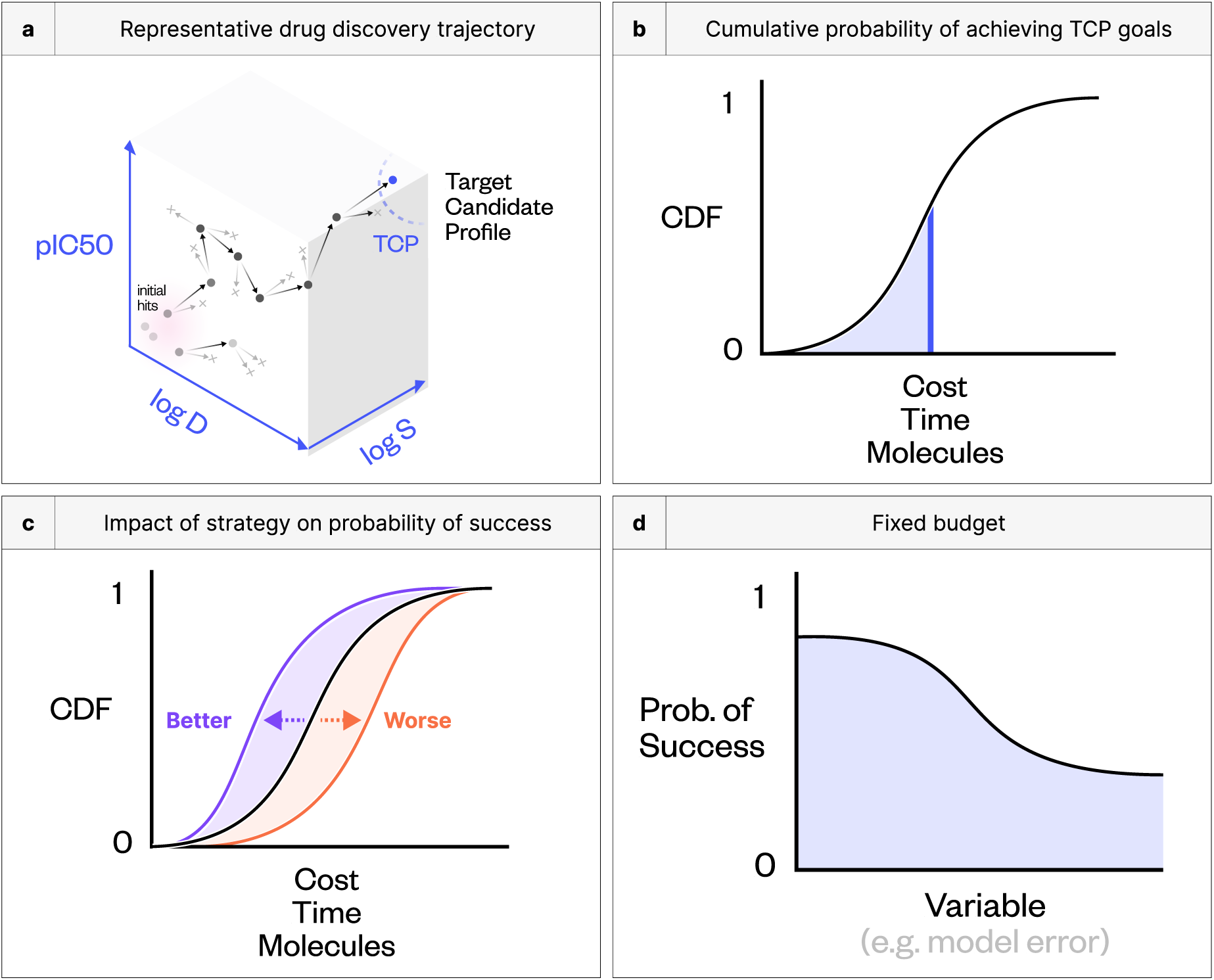
Drug discovery is a stochastic process. **(a)** In drug discovery programs, medicinal chemists must balance multiple, sometimes competing objectives while progressing toward a target candidate profile (TCP). **(b)** Each such trajectory is considered a realization of a stochastic process. The stochastic process can be usefully characterized by cumulative distribution functions (CDF) in terms of important resource or budgetary quantities that may be constrained (monetary, time, or capacity/inventory). **(c)** The statistics of the benefit of different decision-making strategies and technologies can be read off in the shift of the CDF. **(d)** Given a budgetary constraint (e.g., money or time), the probability of successfully achieving TCP goals within the budget constraint can be quantified by examining the behavior of the CDF value as a function of whatever choice is being varied.

How this program unfolds is just one realization of the stochastic process. Running the program multiple times in isolation would allow us to derive statistical measures. For example, a one-dimensional cumulative distribution function (CDF) can describe the probability of achieving the program goals as a function of time, cost, or the number of molecules synthesized in a program (**Figure 1B**). This statistical perspective can usefully characterize the impact of some chosen strategy, model, or technology on the efficiency of reaching the goal. A leftward shift in the CDF signals that the same resources will achieve a higher success rate, or that the expected cost of success is reduced (**Figure 1C**); assuming a fixed budget, the success rate can also be related directly to experimental parameters such as model error (**Figure 1D**).

### DrugGym is a framework for the econometrics of drug discovery decisions

We present DrugGym, a framework for modeling drug discovery as a realistic stochastic process (**Figure 2**). As in a real hit-to-lead campaign, we initialize experiments with hits and iterate through standard design-make-test-analyze (DMTA) cycles until the specified target candidate profile (TCP) goal is reached [53].

**Figure 2.**
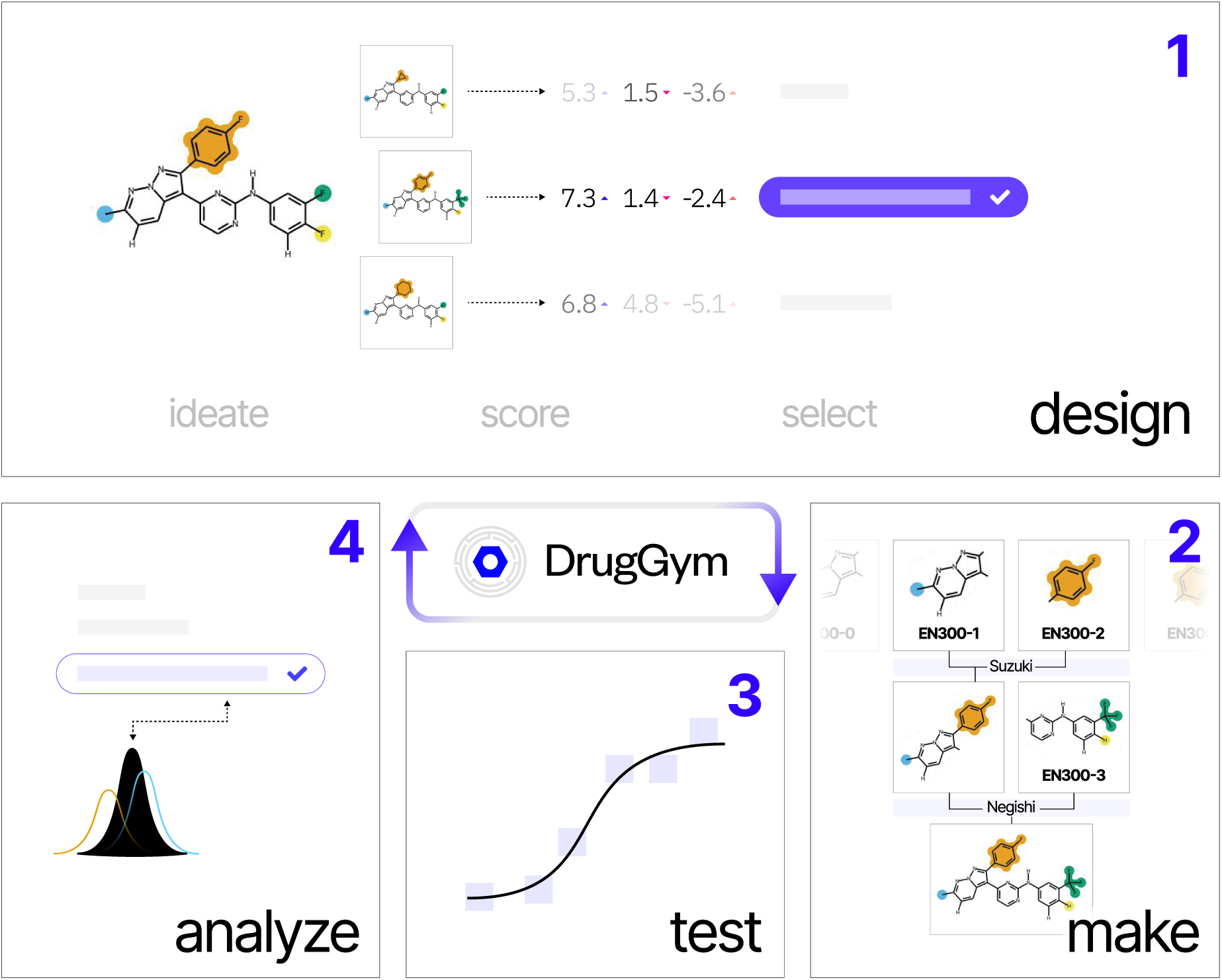
Introducing DrugGym: a model of stochastic drug discovery. Drug discovery proceeds through many design-make-test-analyze (DMTA) cycles, represented in DrugGym by analogous routines. In **design** steps, given assay data about a starting library of molecular hits, molecules are selected for ideating analogs using standard building blocks and robust organic synthesis reactions. Analogs are then scored with surrogate models and selected for synthesis. In the **make** step, designs are synthesized according to their precomputed synthesis route. During the **test** step, molecule properties are measured. In the **analyze** step, available information is used to improve selection in the next cycle.

Runtime is a major concern in how DrugGym was developed. We must perform thousands of simulations to draw meaningful quantitative conclusions, and a single experiment can involve identifying and scoring dozens of synthetically feasible analogs for thousands of compounds. Accordingly, we adopt parallelizable methods with low computational complexity. These can scale to hundreds of compounds simultaneously: milliseconds to identify analogs and seconds to score all of them (**Supplementary Figure 18**). This efficiency allows running 100 large-scale experiments in minutes.

DrugGym models the DMTA cycle with modular implementations so that interventions can be compared within each step for their impacts on success rates and cost-effectiveness:

#### Design

Given starting hits as inspiration, we follow the standard paradigm of generating a number of analogs (*ideation*), scoring these analogs using predictive models (*scoring*), and selecting molecules to be synthesized and assayed (*selection*).

##### Ideation

Given assay data about these hits, high-performing molecules are first selected for ideating synthetically tractable analogs. DrugGym permits ideation strategies to be defined based on synthetic capabilities. Here, we use a simple method that is aware of synthetic routes, substituting synthetic building blocks in the final step(s) to enumerate a number of analogs of each hit. In this way, a vast chemical space (10^21^ molecules) can be rapidly enumerated by combining a large commercially available building blocks catalog (e.g., from Enamine [54]) with a standard reaction repertoire [55].

##### Scoring

We provide a module for scoring ideated designs with predictive models (which may be trained online), although not all human design teams use them. To model the assay process, analogs are scored using realistic computational surrogates with the characteristics of real assay measurements, which DrugGym calls *Oracles* (described in *Test* below).

Modern drug hunters can scrutinize compounds with predictive models before they are ever synthesized and tested. By definition, these models are only approximations. In principle, a set of learnable surrogate models could be pretrained on external data and fine-tuned online (during the *Analyze* step) to model the process of learning from data during the discovery process. Here, our initial goal is to model the impact of unbiased predictive models with varying degrees of inaccuracy. We simulate this characteristic by introducing NoisyOracles, which sample a Gaussian distribution centered on the “true” Oracle value.

##### Selection

DrugGym implements several strategies for selecting scored designs for synthesis and assays. It is sensible to use all available information to select candidates, preferring Oracle values if available, followed by those of NoisyOracles. Yet these objectives are numerous, non-commensurable (different units or scales of measurement), and conflicting (improving one objective may cause another to deteriorate). No universal solution is known for optimizations of this kind [62, 63].

We assess progress toward our TCP goals with a composite *utility function* that maps proximity to each of our objectives into a shared, one-dimensional range. We choose a function that returns a utility for each objective according to its acceptable and ideal bounds in the TCP, selected here to reflect values typical of approved oral drugs (**Figure 3C, 3A**) [64–67]. We desire a single function that saturates when all objectives are satisfied, yet penalizes poor scores without limit. Our utility function maps values within the ideal range to an objective of 1, implying that anything in this range is equally prized. Values observed in acceptable bounds are linearly interpolated between 0 and 1. Those out of bounds receive a quadratic penalty ranging from 0 to ∞ (**Figure 3C, 3D**) [68].

**Figure 3.**
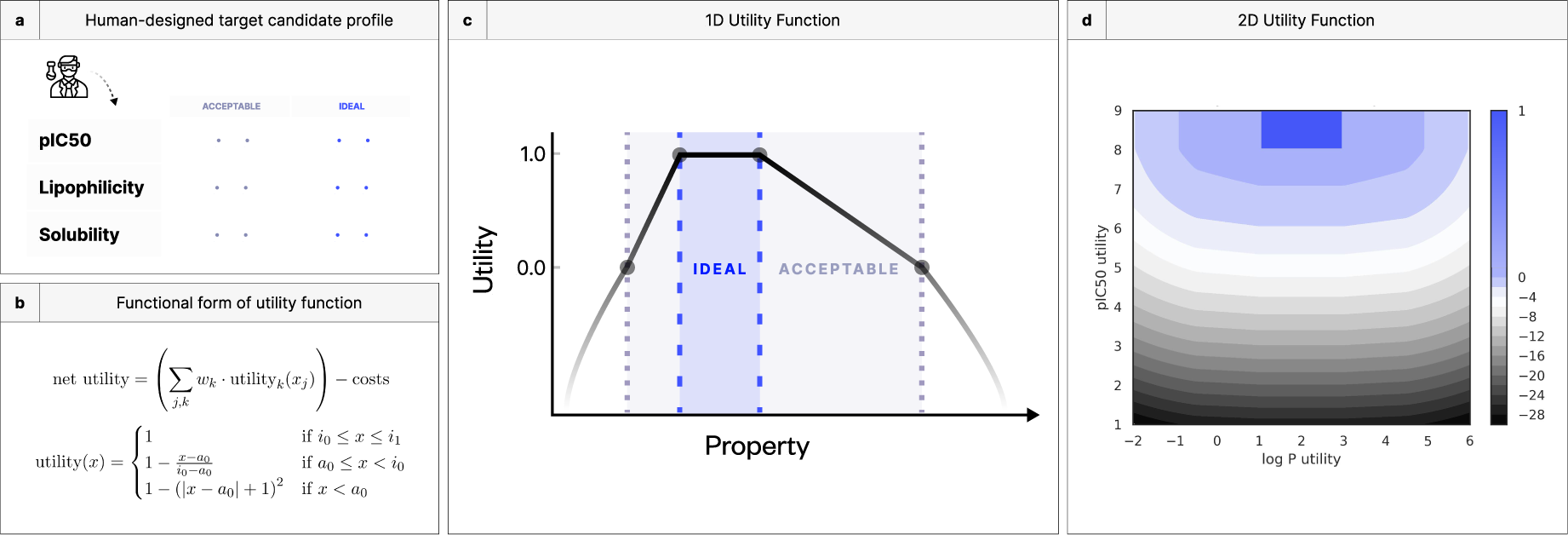
Utility unifies multi-objective problems into well-behaved optimizations. **(a)** Decisions made in every round of a DMTA cycle produce both progress toward the TCP goals and various costs (monetary, time, capacity, etc.). Different projects may value trading off expected progress and cost in different ways. **(b)** We encode the combination of progress and costs into a *utility function* that takes a simple form, inspired by the penalty method: “acceptable” and “ideal” thresholds for each objective can be used to transform arbitrary real-valued objectives into (−∞, 0, 1] bounds. **(c, d)** One-dimensional and two-dimensional illustrations of the standard utility function.

Even with molecule scores in the same dynamic range, it is unlikely that one molecule bests all others across all objectives [29, 69, 70]. We introduce a *selection policy* that can contend with utility trade-offs, in which groups of molecules are non-dominated by each other (the Pareto front).

We first implement a simple policy intended to model the decision-making process of human medicinal chemists. We address conflicts with a two-step procedure. First, we use a genetic algorithm to partition the data into successive fronts of non-dominated molecules [71]. Second, we assign scalarized scores within fronts based on a weighted average of objectives (see **Detailed Methods**).

The selection policy prescribes an optimal ordering of molecules for which predictions and measurements have been made. Recalling that the predictions are imperfect, we can employ reinforcement learning for the final selection [72], balancing exploiting informative predictions with the potential of unexplored chemical space. We use 𝜖-greedy, a simple and competitive baseline that has been applied to many problems [73]. This method selects a molecule at random with probability 𝜖, otherwise defaulting to the policy prescription.

#### Make

By construction, no retrosynthesis planning is necessary, since the ideation step enumerates products using forward synthesis routes from synthetic building blocks. For our virtual experiment, “synthesizing” is assigning an annotation to molecules selected from the yet-unmade collection.

#### Test

A few of the synthesized molecules are chosen to be tested using noiseless Oracles. These include affinity for the ABL1 kinase, estimated using rigid-body docking [56]. We chose this objective because, unlike neural surrogates of affinity [74], docking exhibits two essential features of real affinity measurements: (1) structure-activity relationships between related compounds and (2) activity cliffs, where a single atom change could make or break potency [45]. For the settings we use here, the reproducibility of docking scores mirrors the intra-assay variability of real experimental data collected by the same laboratory [57, 75] (**Supplementary Figure 13**). We score lipophilicity using Crippen’s method [58] and solubility using a gradient boosted tree trained on solubility measurements [61]. These ADME properties are often used to demarcate molecules with the potential to be orally bioavailable [76]. They are also challenging to optimize due to their strongly negative relationship [67].

#### Analyze

As measurements are recorded, they supplant predicted scores and alter selections in future rounds. The goal is to select new designs with potential but avoid neglecting strong molecules that have already been measured. Yet a naive comparison of the two “crowds out” the latter in selected batches, since high predicted scores are overestimates, on average. The overestimation bias, also reported in real programs [16], is most severe for the noisiest scores. To level the comparison, we regress out the bias using complete prediction-measurement pairs (see **Detailed Methods**). Every measurement of a molecule is associated with a cost, so that the predefined budget depletes as progress is made toward TCP objectives. Success is determined by a race of these two variables: successful programs will achieve their objectives before the budget is depleted, while unsuccessful programs will not.

### DrugGym experiments can assess the impact of ideation strategies, batch size, model error, and search methods

Below, we show how we can use DrugGym to estimate program success rates against a representative TCP objective under fixed budgets. Similarly, we show how we can assess the expected costs incurred to achieve the TCP goals, as well as the probability distribution of those costs. We use these metrics to explore the design space of discovery campaigns, varying choices for synthesis, batch size, and search policy. We also investigate consequences of model error, finding that a reduction in affinity model inaccuracy from 2 to 0.5 pIC50 units could improve success rates tenfold. Our work demonstrates a realistic, multi-objective, cost-aware, extendable testbed for applying computational methods to the hit-to-lead phase of drug discovery.

DrugGym, extensive documentation, usage examples, and scripts for reproducing our experiments are publicly available under MIT license at www.drug-gym.org.

## Results

We set a target candidate profile (TCP) objective with affinity, lipophilicity, and solubility goals (**Table 1**, **Figure 4A**) for a hypothetical hit-to-lead phase of a discovery program against ABL1, a kinase dysregulated in cancers such as chronic myelogenous leukemia [77]. Starting with hits from a fragment screening library [78], we step through iterations with DrugGym until a molecule is found that satisfies all TCP objectives. We repeat the experiment to estimate probabilities of success and cost statistics.

**Figure 4.**
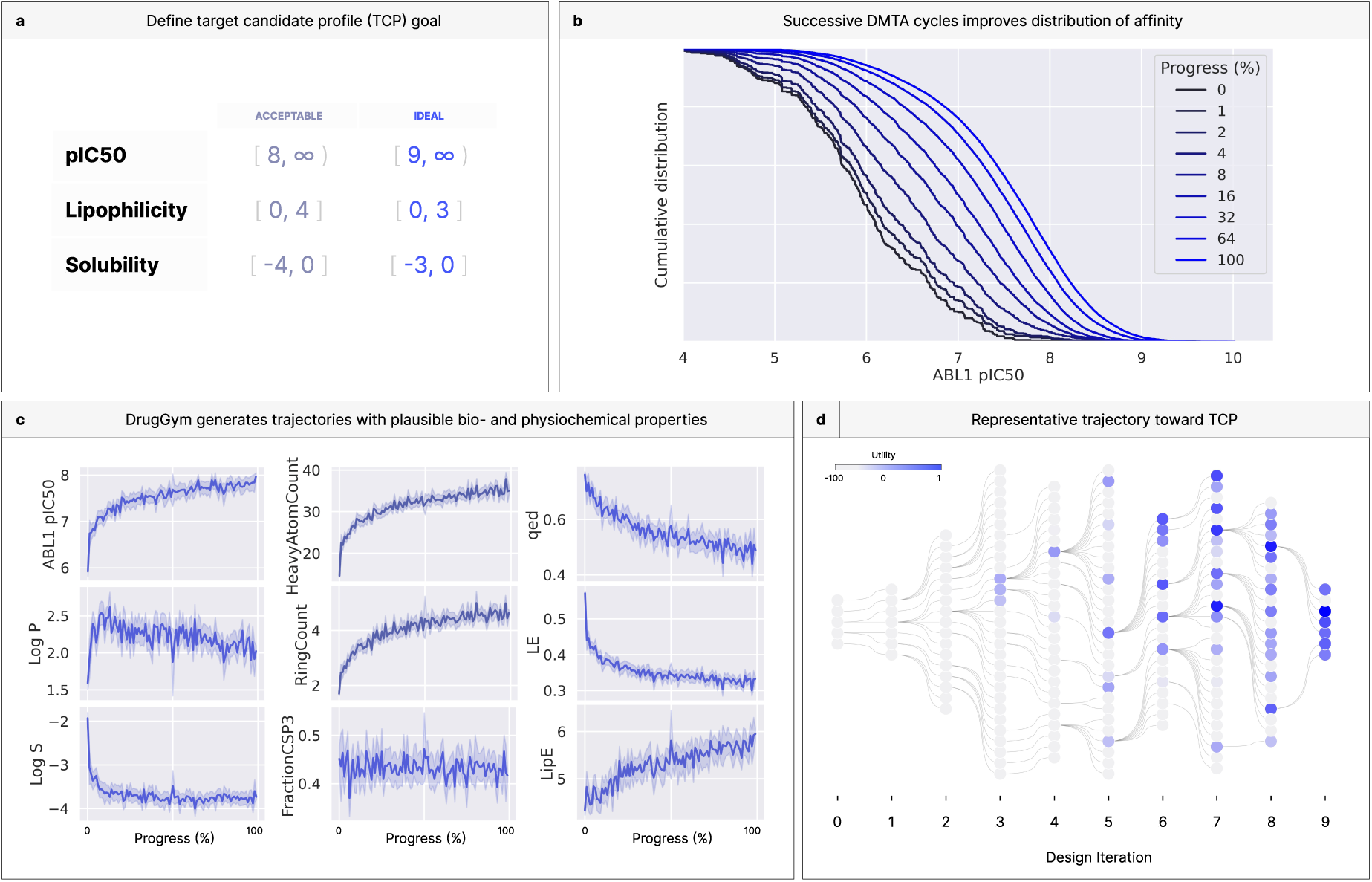
DrugGym simulates discovery trajectories that broadly recapitulate properties of real early-stage drug discovery programs. **(a)** Given a TCP of a simplified hit-to-lead phase (including target affinity, lipophilicity, and solubility goals), we simulate the hit-to-lead progression from hits to sub-micromolar leads with progressable ADME. **(b)** Progress toward the TCP-satisfying lead substantially pushes out the cumulative distribution of ABL1 pIC50. **(c)** Time-dependent averages of chemical properties indicative of drug-likeness averaged over many realizations of the discovery program. The first column represents the TCP objectives; the second represents key indicators of drug-like structures; the third presents typical composite metrics across several objectives (acronyms: FractionCSP3: the ratio of sp hybridized carbons over total carbons; QED: quantitative estimation of drug-likeness; LE: ligand efficiency, pIC50 divided by the number of heavy atoms; LipE: lipophilic efficiency, difference of pIC50 and Log P). These metrics reflect that identified molecules exhibit average metrics generally considered to be drug-like, even while improving average affinity by two orders of magnitude. **(d)** An example of a simulated discovery campaign that reaches the TCP objectives. Edges from left-to-right indicate that one molecule served as inspiration for the next. Design iterations indicate the lineage of synthesized molecules ideated from previous hits, not the total of DMTA cycles elapsed (which may be larger than the number of design iterations for that molecule series).

### Simulated discovery campaigns exhibit plausible medicinal chemistry progressions

Our initial goal with DrugGym is to simulate how a human medicinal chemistry team might ideate and select molecules for synthesis in a real drug discovery program. Due to measurement error, stochasticity in decision-making, and the varying quality of starting hits, programs complete at different times. To make an appropriate comparison across compounds at the beginning, middle, and end of campaigns, we create a normalized measure of campaign progress. Following Beckers et al. [79], we measure progress as the percentage of compounds that have been “made” in the simulation. We use progress percentage to study longitudinal impacts of different ideation and selection strategies.

Aggregating across repeated trials, we show that improvements in the distribution of affinity are strongest in the first third of the discovery programs. After that threshold, each doubling of progress has less effect (**Figure 4B**). We also look at the impact on the expectation of ABL1 pIC50. In **Figure 4C** (top left panel), we see a logarithmic rise in average pIC50, with a fast early phase that continues to rise, albeit increasingly slowly. This shift in the curve is associated with changes in the behaviors of the other TCP objectives (first column, second and third rows). Here, average log P increases initially, before beginning to fall just as affinity ceases its fast phase. In other words, as compounds are identified with affinity in a more acceptable range, they receive a comparatively smaller reward for marginal improvement. This is in line with the design of the utility function. After this point, molecules that can deliver comparable affinity with improvements on other objectives will be rewarded.

On the other hand, average solubility declines precipitously in the early phase. Its trajectory is almost a mirror image of affinity, except that it does halt its downward slide around halfway through the simulation. These results are also consistent with the strong negative correlation we observe between log P and log S (**Supplemental Figure 14**). In effect, the optimizer chooses its battles, finding compounds that improve on two objectives (pIC50 and log P) while holding at bay the negative penalties associated with that choice (decreased log S). After the earliest phase of the optimization, average log P improves by half a log unit and pIC50 improves by almost a full log unit, while log S remains roughly constant. That it stops at log S = −4 is not a coincidence: this is the boundary of the acceptable range that we set in the TCP. The utility function heavily penalizes compounds that fall outside of this threshold. Simulation outcomes are likely to be highly influenced by the design of the TCP, but the degree of impact in our results is unexplored here.

### Simulated discovery campaigns mirror the statistics of real drug discovery programs

Although lacking data for individual compounds, the Novartis series [79] is the largest publicly available study of drug campaign aggregate statistics. We compare our synthetic trajectories against these real programs on several key metrics. Specifically, we assess measures of drug-likeness that include the number of heavy atoms, number of aromatic or aliphatic rings, and fraction Csp3, the fraction of carbons in the molecule that are sp^3^-hybridized (**Figure 4C**, second column). The latter is considered to be a surrogate for three-dimensionality, which is usually associated with stronger ADMET properties [80].

The average number of heavy atoms in the compounds of the Novartis series starts above 30 and rises to around 34. In our simulations, average heavy atom count begins well below 20, before finishing around 35. The large difference in the early part of these trajectories illustrates the difficulty in this comparison, which is that the starting points are likely different chemical matter (ours are fragment hits [81]; theirs may be from a range of sources). Since many properties are downstream of heavy atom count, the most apt comparison might begin after the first third of each campaign, when the heavy atom count crosses 30.

We observe that ring count more than doubles over the course of simulation (**Figure 4C**, second column, second row). However, the story changes when normalizing the number of rings by heavy atom count, following the Novartis series (**Supplementary Figure 15**). Our compounds have normalized ring counts in a similar range (both series converge to ∼133 rings per 1000 atoms). The difference lies, again, in the starting point. They begin their series above this level and drift toward it. In contrast, our simulations begin at around 115 rings per 1000 atoms and rise steadily before equalizing.

Fraction Csp3 exhibits the most notable difference in outcomes, since the average of compounds in the Novartis series rises from <0.30 to about 0.34. The compounds in our campaign have a much higher fraction, hovering throughout between 0.4 and 0.45. A higher fraction is better, since it correlates with solubility and predicts lower CYP450 inhibition [82]. Interestingly, despite large swings in compound size, affinity, lipophilicity and solubility, the fraction Csp3 is virtually flat from start to end. Like ring count, this should be a function of the starting points (hits), accessible chemical space (via available building blocks and reaction repertoire), and selection dynamics.

Composite metrics such as QED [38], ligand efficiency [83], and lipophilic efficiency [83] can be used to relate together several individual molecule descriptors. These reflect that the drug-likeness of our compounds start favorably (QED >0.75) and fall to a region still considered high-quality (>0.5) [82]. Efficiency measures evaluate the cost paid in terms of molecule size or unwanted lipophilicity for increased affinity. This is especially important when entering the lead optimization phase, since it is nontrivial to rescue an inefficient scaffold by decorating its periphery, and scaffold hopping in late optimization is challenging [83]. Our simulations demonstrate that ligand efficiency decreases markedly from its initial posture, but that, as with QED, it converges to an acceptable region (>0.3) [82]. Lipophilic efficiency is perhaps the biggest winner of our simulations, rising almost monotonically from 4.5 to nearly 6 units, on average. The best molecules we observed had >8.5 lipophilic efficiency (**Figure 6**).

We report additional measures of drug-likeness in **Supplementary Figure 15**. These indicate that the average compound structure starts out very drug-like and becomes slightly less so over the course of the discovery campaign, confirming the observation from QED. There are two caveats: first, the averages are still within the range of typical rules of thumb for drug-likeness [59]; second, we did not optimize for any structural objectives in our simulations. The fact that resulting compounds are already drug-like suggests that significant performance gains are possible by explicitly considering these metrics during selection.

### DrugGym enables detailed examination of the lineage of compounds that achieve TCP goals

In **Figure 4D**, we visualize a single discovery campaign in terms of direct inspiration of each compound that is ideated, synthesized, and tested. That is, from five starting hits, edges denote that the compound on the left was used in the Design step to ideate the molecule on the right. Examples include the replacement of a reactant in the parent compound with a similar building block or growing the fragment by including the entire parent compound as a reactant. The panel is colored by utility scores. What is most evident from this rendering is a “jackpot” effect, in which “winning” compounds inspire the vast majority of compounds downstream. This effect validates that our simulation is finding SAR and exploiting it. It also implies that the performance of descendents may be informative of the quality of the ancestor for subsequent selections. Importantly, the choice of locally suboptimal molecules (as selected using our 𝜖-greedy selection procedure) can bear dividends that may only be apparent after several design iterations. For example, in iteration 5, the yet-highest utility compound was identified as a descendent of a molecule that performed poorly. Under a purely greedy selection strategy, that compound might never have been found, or much greater cost and time would have been expended. That is not to say that these “suboptimal” selections always work; they often do not (as in the selection connecting the bottom of iteration 6 to the bottom of iteration 7). As we note in the **Discussion**, machine-guided selection procedures may be employed to balance these concerns.

### DrugGym samples reasonable chemical structures and synthetic routes

Using the same notion of progress (percentage of molecules made so far), we gathered a random sample of compounds from different phases of simulated campaigns (**Figure 5A**). We stratify the compounds by their progress-class, seeking qualitative differences across design cycles. This sample, though sparse, demonstrates that compound size does not monotonically increase across DMTA cycles, as one might expect. Some of the largest compounds are found at 10% and 40% progress; one of the smallest compounds is at 10% progress. Since every DMTA cycle involves a fragment growth phase, compound size will increase with high probability in the absence of selective pressure. This validates our observation in the previous section that selecting for our TCP effectively controls structural parameters in addition to the functional ones it was designed for. Above a certain size of molecule (around heavy atom count 30), affinity gains are marginal while solubility plunges (**Supplementary Figure 14**).

**Figure 5.**
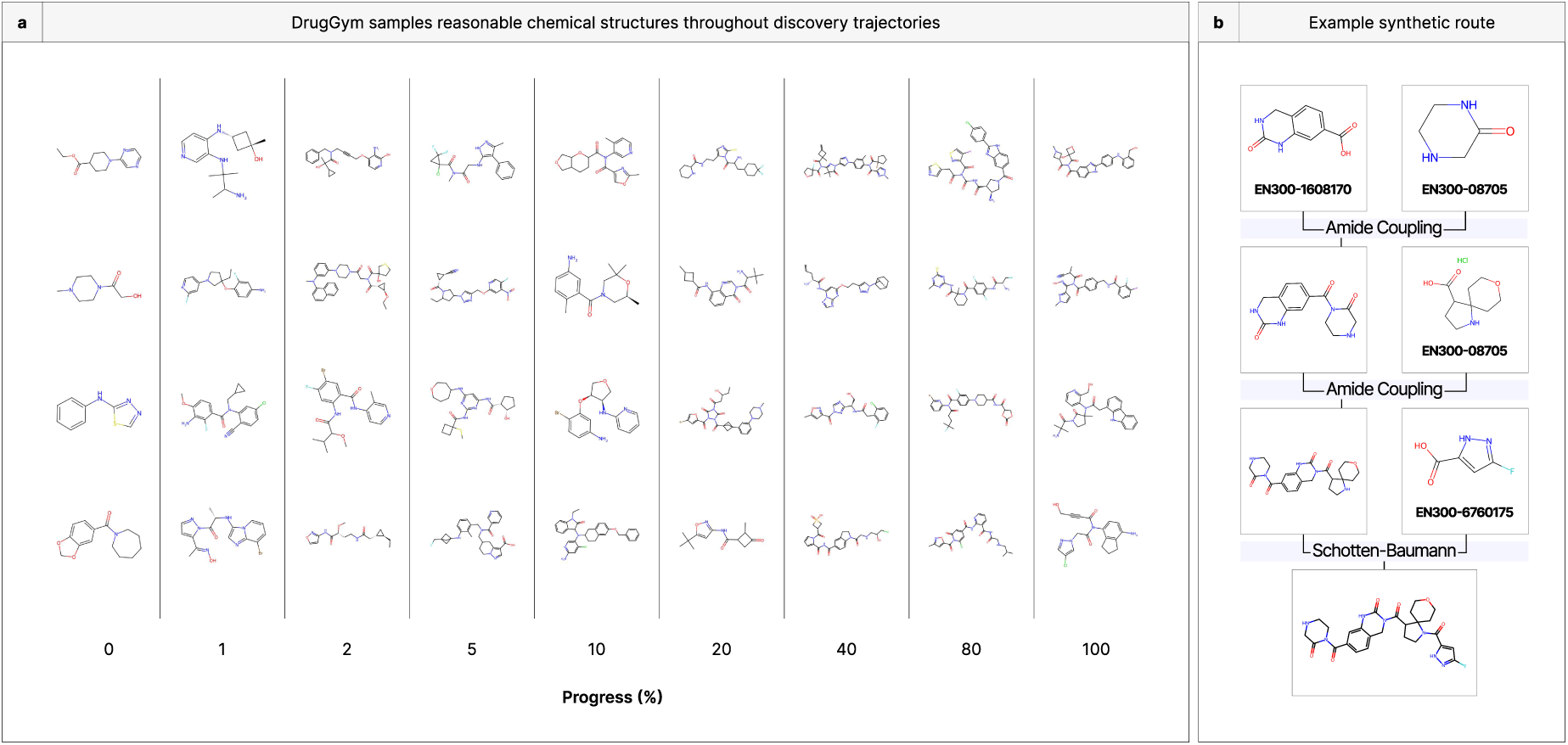
DrugGym samples reasonable and synthetically accessible chemical structures. **(a)** Since programs finish at different times, we compute a composite measure of progress. We sampled molecules at random from these different slices of progression from hit to lead. The resulting compounds exhibit a high degree of diversity of size and structure. **(b)** An example of a synthetic route for a compound satisfying the TCP objectives, including unique identifiers for bulk order from Enamine. DrugGym provides a complete synthetic route for every compound sampled.

We also observe that the generated compounds are diverse. A wide range of structural motifs and atom types are employed, and the simulation avoids retreading old chemical space. This is partially by design, as the Designer module maintains a cache to avoid repetitions.

The designs include liberal use of atoms that have traditionally been less common in approved small molecule drugs, particularly halogens [84]. In our sampled molecules, every progress cross-section after the starting hits includes at least one halogenated compound. Although we have intentionally selected a reaction repertoire and building blocks that are exceedingly popular in the medicinal chemistry toolbox, our selection process is not influenced by prior dogmas (e.g., atom choices) [85]. The default selection routine in DrugGym does not explicitly consider structure, relying instead on scores from surrogate models and measurements from Oracle assays. As a result, our system could make unbiased choices that would be out of the ordinary for a human chemist. Follow-on studies could test encodings of traditional heuristics for structural selection, such as PAINS filters [86].

Of course, the total population numbers in the millions of compounds, so a snapshot of 36 representative molecules is very far from a complete picture. Additional designs and the complete library are available at www.drug-gym.org.

One representative synthetic route is shown in **Figure 5B**. This is a 3-step reaction that is typical of molecules identified in DrugGym: it uses simple, robust reactions and building block that can be readily ordered. Hence, even compound suggestions that are formed in multi-step reactions are very synthetically tractable. Despite this, the size of possible compounds is enormous (10^21^, **Supplementary Figure 16**).

Taking the campaign that yielded the molecule in the previous example, we found the sequence of molecules with the highest utility observed at the time they were made (**Figure 6**). These are all within a somewhat related lineage, since most of them are descended from the same ancestor; however, one does not directly inspire the next. This sequence demonstrates that structural changes across DMTA cycles can be quite large. There is little exact overlap between the first hit and subsequent molecules, although the ester pattern is generally repeated. The trajectory also illustrates that successful compounds do not necessarily grow in one direction. Instead, they can wrap in different directions, as in cycles 10 to 11. Critically, many cycles can pass before a new best compound is observed. This confounds rigid early-stopping rules and points to the need for new methods for inferring signals of futility during campaigns. The same phenomenon also demonstrates that very similar structures can be maintained across cycles, such as from cycle 3 to 10, in which the molecule is mostly intact but for a change around the heterocycle on its right flank. In terms of the TCP objectives, this trajectory is a microcosm of the global trends, in that affinity and lipophilicity were each improved by more than two log units, yet solubility remained stubbornly constant.

**Figure 6.**
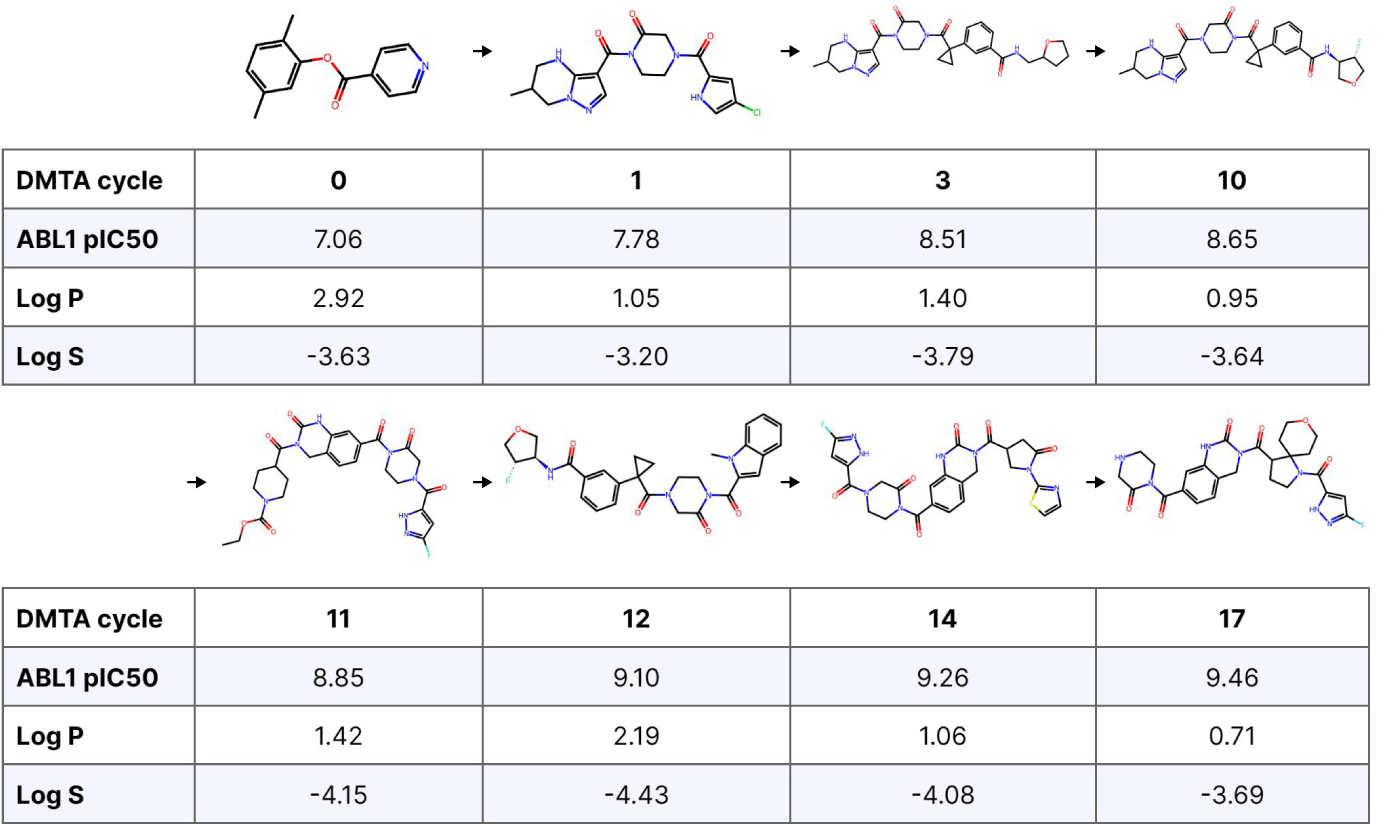
Progression of compounds with the highest utility across design cycles in a simulated discovery campaign. Arrows only indicate program progression, not direct inspiration for the subsequent design.

For our final case study, we examine the lineage of a single molecule that satisfies our TCP goals (**Figure 7**). This lineage traces the arc of inspiration from the final molecule back to the starting hit. Although the chemical moves are not as large as in the previous example, this was a much longer optimization, running for 49 cycles. Nine cycles elapse between compounds 4 and 5 alone. Here, lipophilic efficiency was not much improved by optimization. Much of the improved affinity may thus be explained by increased hydrophobicity.

**Figure 7.**
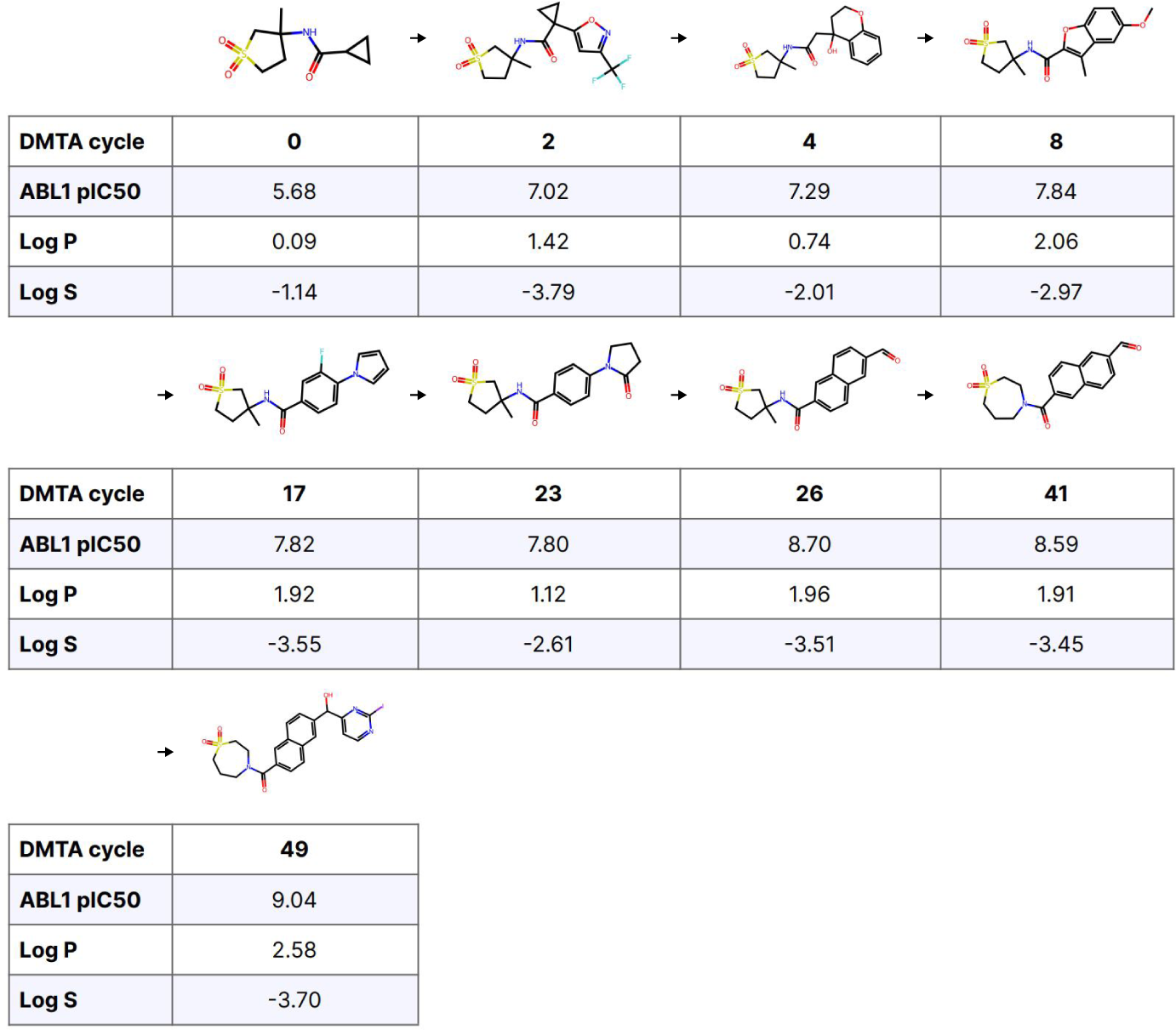
Molecule lineage for a single compound satisfying TCP objectives. Here, arrows indicate direct inspiration for the following compound in the sequence during the ideation step. In the synthesis of that compound, the parent compound participated in-whole or in-part and was used to identify similar building blocks.

Overall, the compounds identified in our simulations are drug-like and can be made using realistic synthesis plans. The longitudinal dynamics also conform to aggregated metrics of real-world drug discovery programs, such as the Novartis series. Deeper study of these dynamics may validate new optimization strategies in the hit-to-lead phase.

### DrugGym finds synthetic strategies for hit expansion against a drug target

Based on trajectories like the ones outlined above, we sought to relate the creativity of chemical moves during the Design phase and resulting program outcomes. By default, our Design phase has two stages: replacement and growth. The growth phase involves selecting a random building block (subject to size constraints) to join the parent compound in a compatible reaction. In this case, we do not have a well-defined notion of creativity.

During replacement, a reactant from the synthetic route is replaced with a member of a building block library, chosen according to fingerprint similarity (**Figure 8A**). Initially, all building blocks are ranked in this way, and the ranking is reweighted according to a Boltzmann sampling procedure (see **Detailed Methods**). The “temperature” of this procedure determines the degree of faithfulness to the initial ranking. A temperature close to 0 will be maximally conservative, while a temperature above 0.2 may deviate significantly; a temperature of 1.0 is akin to random selection. This temperature serves as the first parameter controlling the degree of creativity. The second parameter is simply the number of reactants replaced according to this procedure (“𝑛-replacement”). This is crucial, since maximizing temperature for only one reactant will reach a ceiling on the global dissimilarity that can be reached compared to the parent molecule (**Figure 12B**). For this reason, the most dissimilar descendents are made when all reactants are replaced (all-replacement) according to a random search (infinite temperature). In addition, the impact of temperature is amplified as more reactants are replaced. This may be because if several reactants are selected above a certain temperature, they may introduce new reactive atom pairings, yielding grossly dissimilar products. In contrast, the most conservative configuration would be 1-replacement with 0.0 temperature.

**Figure 8.**
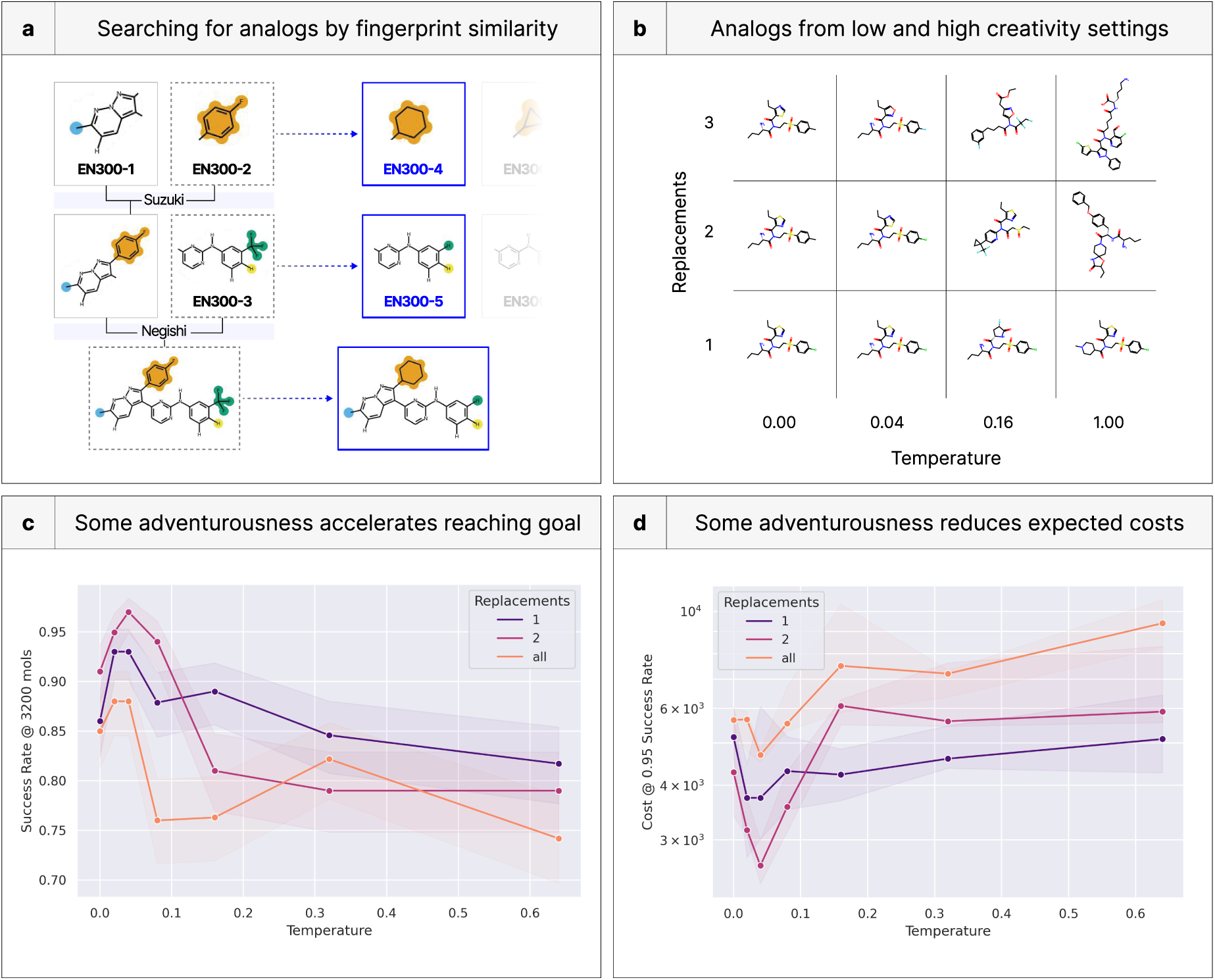
DrugGym discovers optimal ideation strategies for hit expansion against a real-world drug target. **(a)** As a surrogate for the creativity of a medicinal chemist, we enumerate building blocks according to fingerprint similarity to reactants of the original hit. In this setting, creativity can be controlled with a temperature, parameterizing a Boltzmann resampling procedure on the building blocks. **(b)** Representative molecules from low- and high-creativity settings. **(c)** Success rate as a function of the temperature parameter and the number of reactions replaced in analogs versus the parent compound, given a fixed budget of 3200 compounds made. Success rate can be very sensitive to the choice of design parameters, impacting the efficiency of overall discovery. Maximizing creativity makes it hard to ascend the optimization space, yet minimal creativity is not optimal, since it may be prone to becoming trapped in local minima. **(d)** Expected cost for a given minimal success rate of 0.95 for different numbers of reactants replaced. The expected cost can depend strongly on ideation creativity.

### A balance of originality and continuity in ideation improves success rates and reduces costs

We show that some chemical creativity raises success rates for fixed budgets, but that too much adventurousness begins to hinder a successful search (**Figure 8C**). This is especially evident at the lowest temperatures, where the same pattern is observed across all replacement settings: success rates rise from 0.0 temperature before peaking at 0.04. The curves then begin to fall before smoothing out at high temperatures (0.32 and 0.64). These findings validate our prediction that the lowest temperatures could be more prone to getting stuck in local minima. At low temperatures, 2-replacement is favored, with a top success rate above 0.95 at temperature 0.04 given the fixed budget. Then, at higher temperatures, this setting underperforms 1-replacement. This may be because of the “amplification” effect we describe above. The underperformance of the highest creative settings follows basic medicinal chemistry principles, since large changes to scaffolds could disrupt binding modes and undermine steady progress on ADME properties like solubility and lipophilicity. No temperature dominates the others across all replacement settings, suggesting that static creativity could be outperformed by dynamic creativity. For example, ideation could be more adventurous early-on before leveling off in late optimization.

The change in success rate translates into trends in expected cost (**Figure 8D**). We find that the most successful creativity setting (2-replacement with 0.04 temperature) also has the lowest cost in terms of number of compounds made before reaching the TCP goal, at 2618. The highest cost was 9402 compounds, for all- replacement with 0.64 temperature. Given that all other settings were held constant, it is striking that costs can be reduced fourfold by smarter analog ideation strategies. As with the success rate, 1-replacement passes 2-replacement in performance between 0.08 and 0.16 temperatures.

### Batch size drives trade-off of time- and cost-effectiveness

The size of each batch has high practical significance: program leaders may place differently sized synthesis and assay orders depending on the utility they attach to funding and to time to market. We characterize the impacts of fixed batch sizes on program outcomes and expected costs to reach TCP goals (**Figure 9**).

**Figure 9.**
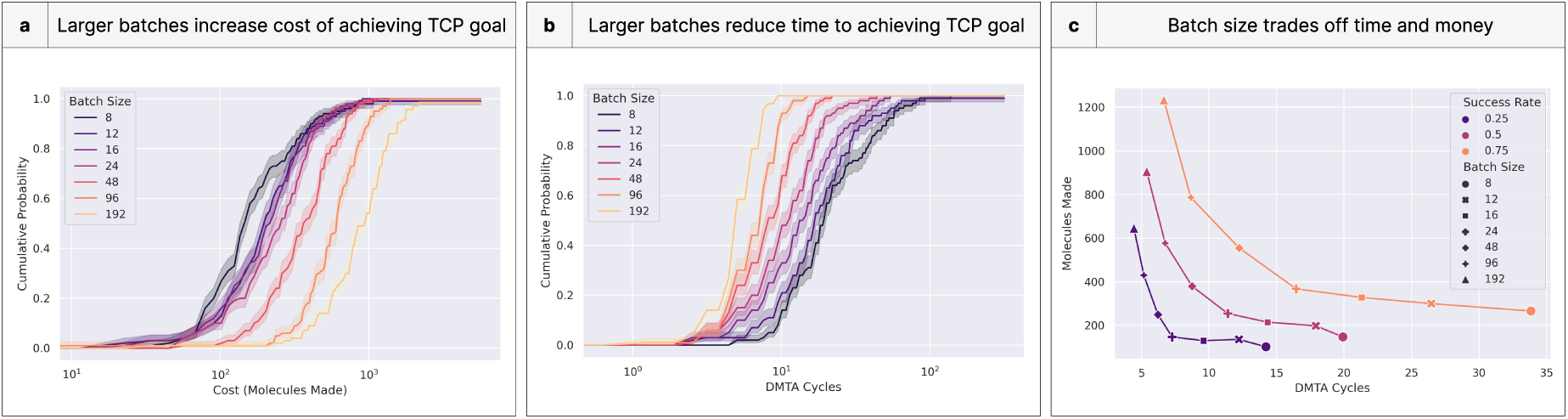
Batch size determines a Pareto frontier of time and monetary efficiency. **(a)** Increasing size of each batch tends to trade off additional monetary cost at the expense of time because there is no opportunity for a surgical search of chemical space. This is akin to a breadth-first search. **(b)** When comparing the DMTA cycle (time cost) rather than the number of molecules made, the larger batch sizes increase the probability of success by searching a larger chemical space in each iteration. **(c)** Batch size trades off one dimension of cumulative distribution function for another (e.g., large batch sizes trade off money for time). This efficiency frontier of cost versus time indicates opportunity for different institutions to value their net utility differently (for example, favoring rapid progress despite exorbitant costs).

The results are not surprising: batch size is correlated with greater costs in terms of molecules made (i.e., money; **Figure 9A**) and inversely correlated to the number of DMTA cycles needed to reach goals (i.e., time; **Figure 9B**). Interestingly, there is little difference in the molecule costs for small batch sizes above a 0.80 success rate (**Figure 9A**). The steepness of the curves in other regions suggests the convergence is not caused by outperformance of the mid-sized batches, but rather the premature flattening of the smallest sizes. This could be an artifact of our simulation, but it also suggests that the most challenging optimizations may benefit from larger batch sizes. Batch size is closely related to breadth-first (large batch) and depth-first (small batch) search strategies, and this result signals that breadth-first search could outperform in such settings.

The primary benefit of larger batches is time savings, since otherwise one could surgically explore chemical space sequentially, one compound after another. The most interesting result from the second panel is that the leftmost curves have not yet saturated (**Figure 9B**). This suggests that relative time-savings may continue to accrue to even larger batch sizes.

By examining expected time and monetary costs to reach our TCP goals, we identify an efficiency curve (**Figure 9C**, lower left is optimal). Across success rates, the story largely reflects that of the previous two panels, in that small batch sizes are more cost-effective, while large ones are more time-effective. Beyond illustrating these tradeoffs, the “knees” of the efficiency curves can be used to identify points of diminishing marginal returns. For example, at the lowest success rate, batch sizes above 24 have diminishing marginal returns on time; those below have diminishing returns on cost. The sharpness of this knee varies by success rate. Arguably, the knee of the curve for 0.75 success rate is 48. Since drug programs may require differing success rate guarantees (depending on size of market, portfolio, and corporate priorities), different batch sizes will be optimal.

Of course, batch size often changes throughout assay cascades, as the earliest batch sizes may be very large, but later optimization has a far lower throughput. The success of different batch sizes may also relate to synthetic capabilities, as a larger batch may gain comparatively more utility from densely sampling a region of chemical space.

### Model error strongly impacts probability of success

As previous authors have emphasized [41], predictive validity can make a large difference in program success rates. If the difference is large enough, even very “inefficient” predictive models can be cost-effective [41]. We analyze this phenomenon in our DrugGym sandbox by adding a layer of Gaussian-distributed model error on top of ground truth, which we derive using our Oracle surrogate models (**Figure 10A**). We experiment with the spread of the error distribution (parameterized by 𝜎) for its effect on success rates and expected costs.

**Figure 10.**
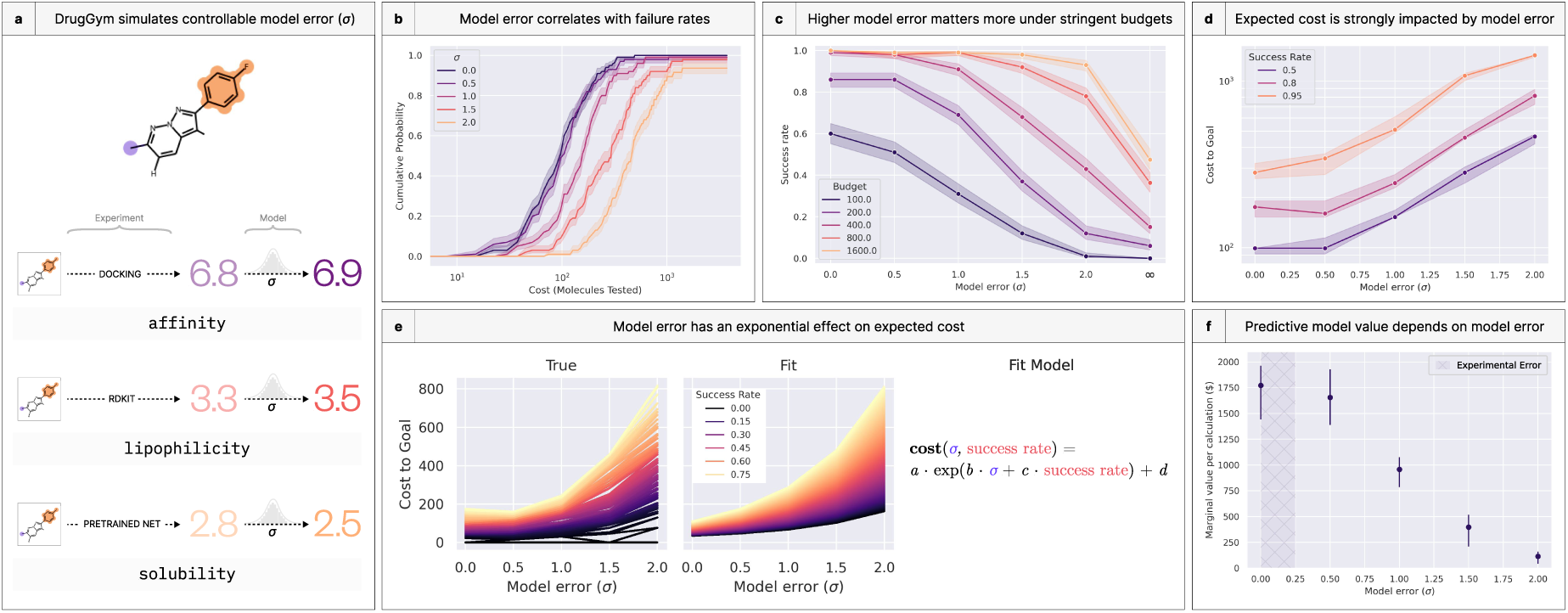
DrugGym simulations demonstrate that model error strongly impacts probability of success. **(a)** The scoring routine in DrugGym contains a set of oracle functions that model properties of interest. We add a Gaussian noise distribution on top of those properties to test the sensitivity of success rates to escalating model error. **(b)** The resulting CDFs are sensitive to error in predictive models. At the lowest sigma, model noise may be eclipsed by intrinsic error in the experiment, abrogating further leftward shift. **(c)** Probability of success falls more steeply when the available budget is lower. The decrease in probability of success is approximately monotonic. **(d)** Expected cost of discovery campaigns rise dramatically with high model error. **(e)** This rise in costs is well-described by an exponential model. **(f)** With very high model error, scoring has no limited effect on selections. By comparing costs of these trials to those with low error, we can quantify the benefit from the scoring models. We illustrate this additional benefit as the median marginal value of each calculation made. This also represents the expected breakeven price of each calculation made. We again note that very low model error still has irreducible experimental error (shaded region), so marginal value may not improve in noiseless settings.

We find that model error is highly correlated with failure rate (**Figure 10B**). The rightward shift in the cost curve is consistently a quarter of an order of magnitude for every 0.5 increase in model error. We examine the functional form of this relationship later. But there is a notable exception at the lowest model error. Here, the rightward shift is slight. This is probably not resulting from a deficiency in the 𝜎 = 0.5 case, but rather a pathology of our “noiseless” experimental data. Depending on the chemical space being scored, our pretrained model of log S has non-negligible error, with a standard deviation of 0.25 to 0.37, and we estimate that our ABL1 pIC50 Oracle has a re-docking standard deviation of 0.15 (**Supplementary Figure 13**). This means that there is an effective floor on the error associated with our experimental settings. On the other hand, the rightward shift at high error shows no signs of abating. We therefore ran an additional experiment with model error set to an extremely large number, far greater than the dynamic range of the objectives. As shorthand, we refer to this as the 𝜎 = ∞ setting. This experiment contextualizes our higher end of model error against its theoretical maximum, which we do in the panel that follows.

In our experiments, the success rate of experiments plummets with high 𝜎, and this is most acute in low-budget regimes (**Figure 10C**, budget refers to the number of molecules that can be made during a program). With a budget of 100 molecules made, the success rate for 𝜎 = 2.0 is just 0.01, compared to 0.51 for 𝜎 = 0.5. That gap narrows to tenfold (0.08 to 0.82) around a budget of 175. Only with a 1600 molecule budget can 𝜎 = 2.0 approach parity. Even so, this is far superior to the infinite error case, in which more than half of programs fail at this budget. These scenarios illustrate that bringing a discovery campaign to completion with high model error is especially costly. The budgets required for high success rates can swell by orders of magnitude for every incremental change in 𝜎 (**Figure 10D**). Again, this trend is attenuated at the lowest levels of model error, so much so that the cost differences are well within the confidence interval, even rising slightly around 0.80 success rate.

We quantify the growth of these cost curves by fitting an exponential model to these data (**Figure 10E**). Our model predicts what the budget must be to achieve a desired success rate as a function of model error (𝜎). Despite its parsimonious form, it is a strong fit (𝑅^2^ = 0.989, MAE = 17.2 molecules). The fit is weakest in regions with the lowest error, where the empirical data are costlier than predicted, and with the lowest success rates, where they are unexpectedly efficient.

Returning to predictive validity [41], we estimate the value of the models themselves, per scoring calculation (**Figure 10F**). To do so, we run the same experiment but without the scoring model. Instead, we make every molecule we ideate, analogous to medicinal chemists choosing which compounds to progress based only on experimental data and chemical intuition. Assuming each compound costs $3000 to make and test [87], we derive an expected monetary cost from the 50th percentile of these trials. We compare this cost to that of the experiments with models in the loop, stratified by model error, reflecting the “savings” gained by using those models. Last, we obtain the per-calculation value by dividing into the number of molecules scored. This quantity also represents the breakeven price for each calculation: above it, use of the scoring model is not expected to be cost-effective. The breakeven value of a model with 𝜎 = 0.5 pIC50 units is $1655.38, 14.5 times that of a 𝜎 = 2.0 model ($114.39). These results quantify that the incentive for developing higher fidelity predictive models is high, especially if the cost of this development is amortized across subsequent calculations. Since the costs of synthesis and testing can easily exceed $3000 per molecule [87], especially in late-tier assay cascades, these estimates should be considered a lower bound.

Beyond these findings, our analysis presents a template for testing new predictive models for impacts on success rates and expected costs in a realistic testbed of drug discovery.

### Scoring models can make up for an impoverished budget

We extended this experiment to quantify gains from a higher ratio of molecules scored versus molecules tested (“scoring ratio”; **Figure 11**). Fixing 𝜎 = 1.0, we find that a higher scoring ratio indeed increases success rates (**Figure 11A**), presumably because it samples more chemical space. The exponential leftward shift in the CDF of successful programs seems to accelerate toward the 20 scoring ratio curve, suggesting that greater gains could come from scoring even more compounds.

**Figure 11.**
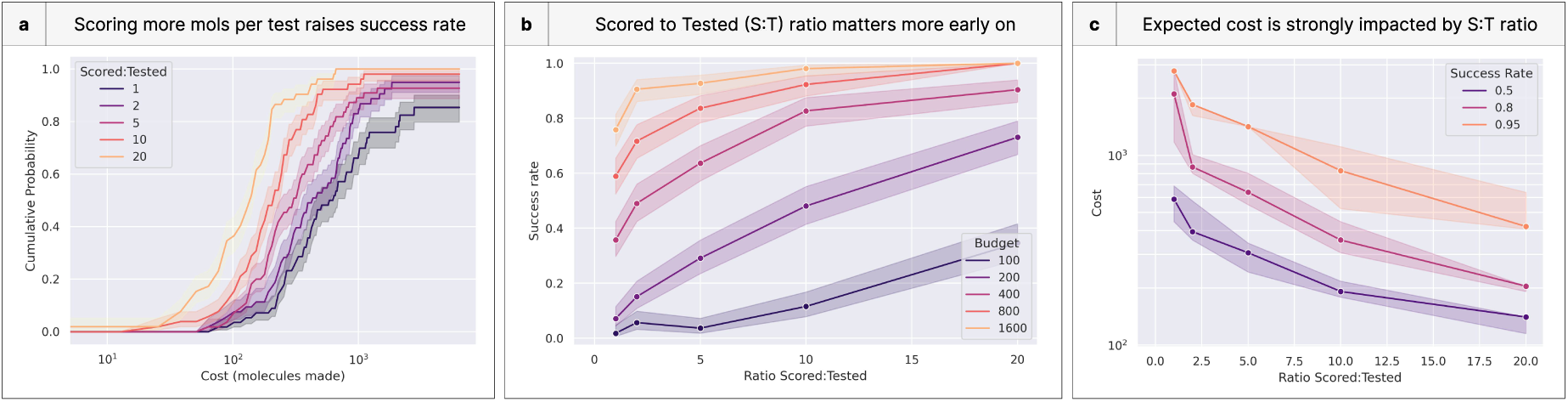
Scoring models are most useful earlier in discovery programs. **(a)** Cumulative distribution of program success for escalating budgets of molecules made. Scoring more molecules prior to testing them substantially improves program efficiency. **(b)** Higher ratios of molecules scored to tested are more impactful in low-budget regimes, where they can raise success rates by nearly an order of magnitude. **(c)** This success rate also translates to the expected costs. While the highest success rates are the costliest to achieve, those costs drop with a greater number of molecules scored.

These effects are most profound early in programs, as scoring many molecules can overcome a small budget (**Figure 11B**). For example, scoring 20 molecules for every one tested raises success rates to 0.73 on a budget of 200 molecules made (compared to a success rate of 0.07 with no such scoring). Correspondingly, the scoring ratio is negatively correlated with expected costs (**Figure 11C**).

## Discussion

### An econometric view of drug discovery is useful for assessing the value of predictive models

While the use of predictive models in drug discovery has a long tradition [88–90], it has been difficult to quantify the value of given predictive models in terms of how much they reduce the time and cost of discovery.

Here, we have demonstrated an end-to-end harness for assessing the value of predictive models, reporting average cost savings in discovery programs for every calculation performed. This arithmetic provides a straightforward estimate of the expected value a predictive model delivers to discovery. Predictive calculations that can be performed at a cost lower than this value will result in overall cost savings, on average. Our approach represents a quantitative rubric for biopharma leaders deciding where and how to invest in models with greater predictive validity.

While our current experiments consider only a single, unbiased notion of inaccuracy (𝜎), DrugGym can be extended beyond this paradigm. Individual predictive models could be evaluated with consideration of their biases [91], multiple levels of fidelity [92, 93], and asymmetric, non-Gaussian error distributions [94]. In addition, predictive models, and the statistical phenomena they capture, are typically co-dependent [95] (**Supplementary Figure 14**). Certain ensembles of predictive models with correlated errors may attain different values. An extreme case would be predictive models that are valuable on their own but totally redundant—their collective predictive validity would then be insufficiently enhanced to outweigh their combined costs. This scenario illustrates that predictive modeling can have significant opportunity costs.

### DrugGym is a convenient sandbox for exploring the econometrics of drug discovery

We developed DrugGym as a convenient framework for exploring the impact of different ideation strategies, predictive models, and search methods on the econometrics of drug discovery. It can be used to track costs (and times) associated with both syntheses and assays of varying complexity and cost, since it was designed to model every step of the Design-Make-Test-Analyze (DMTA) cycle [53] in a general and modular fashion. These costs and durations can be dynamically rendered, with marginal costs rising in later development cycles or as a function of the number of synthesis steps required to make a compound. One can also model the effects of different choices for replications or confirmatory assays, which would reduce uncertainty at greater cost.

### DrugGym provides a platform for exploring decision-making strategies in drug discovery programs

The current form of DrugGym mimics the decision-making strategies of human medicinal chemists in the hit-to-lead stage [53], but more complex decision-making strategies can be explored as well. For example, Bayesian optimization criteria that balance exploration and exploitation may provide value beyond simple “greedy” strategies [69, 96–98]. Methods that look ahead several steps using Monte Carlo Tree Search (MCTS)-like strategies may be able to anticipate “dead-end” actions before they are selected [99]. These methods have performed well in complex open-ended games such as Go [100] and in relevant chemistry applications [11, 12, 21]. Here, they may eschew highly rated chemical spaces that will rapidly plateau, or find that some chemical spaces are more amenable than others for identifying promising molecules. If constraints in synthesis and assay capacity are modeled appropriately, these advanced strategies could account for the opportunity cost of decisions—when making/assaying one compound likely means another cannot be made and assayed—which is very difficult for human teams to assess [101].

### DrugGym can form the basis for exploring reinforcement learning strategies for autonomous drug discovery

DrugGym extends the Gymnasium (formerly Gym) framework [102], a popular choice for implementing reinforcement learning (RL) sandbox environments that attempt to faithfully model real-world tasks. With this foundation, DrugGym can be extended to a fully RL context and used to explore RL policies, enabling decision-making agents to back-track or even stop entirely if further progress is deemed too costly or unlikely. Results could be compared to traditional assay cascades, which gate entry criteria to a set of assays stratified into tiers of increasing costs. We expect that autonomous RL policies may learn more flexible policies that consider both time and cost [103], enabling assays to be performed eagerly, “at-risk.”

### DrugGym can be augmented with learnable predictive models to drive real discovery programs

Predictive models are mimicked in DrugGym by adding random errors to an “Oracle” intended to represent real assays. By replacing these model surrogates with learnable predictive models and using real assay data instead, DrugGym could drive decision-making in real programs [104]. Additional predictive models for more complex ADMET and PK properties involved in real discovery programs can be integrated [105, 106]. The oracles can still usefully evaluate decision-making strategies in a manner that can be applicable to real programs, provided the oracles have the same character as the real assay data (e.g., structure-activity relationships punctuated with activity cliffs [45]). The timing and costs of model training could also be incorporated into DrugGym’s program accounting as part of the Analyze step.

### Current limitations of DrugGym span the DMTA cycle

Although it is a relatively realistic model of the discovery process, DrugGym has several notable discrepancies from real-world design cycles that limit the interpretability of its results. These may be addressed in future evolutions.

#### Ideation strategies in the Design step

Ideation strategies in DrugGym are limited to replacing reactants and adding new ones, and, for the latter, this is without regard to chemical structure. Our current Design step cannot conceive changes that involve simultaneous additions and replacements that could be tuned for similarity to the original compound. Relaxing this constraint, possibly by looking ahead several steps, might enable more nimble chemical moves than are possible with available building blocks. In another sense, our synthetic enumeration is too relaxed. At low ideation temperatures, substituents are often so similar to the originals that the previously reactive atoms are also preserved (see **Detailed Methods**). But we do not enforce this as a hard constraint, and core scaffolds can get disrupted. We have implemented a method for “protecting” non-participating atoms (**Detailed Methods**), but this may not prevent side-products from being enumerated. An additional mitigation could be adding a final similarity threshold after enumeration.

The chemical space that is accessible during ideation is a function of predefined building blocks and reactions. While the cardinality of this space is huge (**Supplementary Figure 16**), these choices could exclude promising classes of molecules [107, 108]. For more complex synthetic pathways, molecules could be ideated with sophisticated retrosynthesis models like AiZynthFinder [18, 109] and Sparrow [110].

#### Limitations of Oracles in modeling reality

Currently, DrugGym models target inhibition with docking scores since these capture key features such as structure-activity relationships and activity cliffs. However, docking scores are highly imperfect surrogates of reality [111, 112]. To aid with compound-level decisions in real-world programs, DrugGym could be equipped with an Oracle model with stronger correlation to experimental target activity. We anticipate extensions of DrugGym to an active learning setting, in which a QSAR model is trained in a loop with experimental data [104, 113]. Additionally, one may seek to include structural objectives that are simpler surrogates for ADMET/PK properties, such as molecular weight, heavy atom count, TPSA, and aromatic ring count [114]. Our present experiments do not consider these objectives (though, as discussed, they often maintain drug-likeness in practice).

#### Improvements in the Analyze step

One of our observations from examining molecule design progressions is a “jackpot” effect, in which some high-performing compounds are far more prevalent in the lineage of chemical designs. The converse can also be inferred: the quantity and quality of descendents may be informative of the fitness of ancestors. This finding could be instrumentalized in the Analyze step by discounting (or rescuing) progenitors’ estimated utility for subsequent selections based on the performance of their progeny.

### DrugGym can be adapted to test new interventions in drug discovery

We have introduced DrugGym, a realistic testbed for assessing interventions across the DMTA cycle of drug discovery programs. As interest in computational drug discovery grows rapidly, it is essential for practitioners to adopt rigorous benchmarks that reflect standard practices of medicinal chemistry and statistics of discovery campaigns. To this end, we have made DrugGym, documentation, usage examples, and scripts for reproducing our experiments, publicly available under MIT license at www.drug-gym.org. Our reusable components are easily adaptable for new predictive models, synthesis strategies, approaches to decision-making, and econometrics analyses.

### Disclaimer

The content is solely the responsibility of the authors and does not necessarily represent the official views of the National Institutes of Health.

### Disclosures

JDC is a current member of the Scientific Advisory Board of OpenEye Scientific Software, Redesign Science, Ventus Therapeutics, and Interline Therapeutics, and has equity interests in Redesign Science and Interline Therapeutics. The Chodera laboratory receives or has received funding from multiple sources, including the National Institutes of Health, the National Science Foundation, the Parker Institute for Cancer Immunotherapy, Relay Therapeutics, Entasis Therapeutics, Silicon Therapeutics, EMD Serono (Merck KGaA), AstraZeneca, Vir Biotechnology, Bayer, XtalPi, Interline Therapeutics, the Molecular Sciences Software Institute, the Starr Cancer Consortium, the Open Force Field Consortium, Cycle for Survival, a Louis V. Gerstner Young Investigator Award, and the Sloan Kettering Institute. A complete funding history for the Chodera lab can be found at http://choderalab.org/funding.

## Funding

Research reported in this publication was supported by the National Institute for General Medical Sciences of the National Institutes of Health under award number R35 GM152017. This research was funded in part through the NIH/NCI Cancer Center Support Grant P30 CA008748.

JDC acknowledges support from the Sloan Kettering Institute. MR acknowledges support from the Tri-Institutional Program in Computational Biology and Medicine and the Sloan Kettering Institute.

## Acknowledgments

The authors would like to thank Theo Karaletsos (**ORCID:** 0000-0002-0296-3092) for valuable discussions on modeling the drug discovery process within a reinforcement learning framework, and both Melissa Boby (**ORCID:** 0000-0003-1920-206X), Jenke Scheen (**ORCID:** 0000-0001-9781-0445), and Ed Griffen (**ORCID:** 0000-0003-0859-554X) for invaluable discussions about the activities of medicinal chemists within DTMA cycles. We thank the scientists of the ASAP Discovery Consortium for fruitful discussions about drug discovery objectives and practices.

The authors are grateful to the MSKCC DigITs and HPC team, especially Jamie Cheong, Lohit Valleru, Svetlana Mazurkova, and Monica Chakradeo for their assistance with high-performance computing resources.

## Appendix: DrugGym

### Detailed methods

#### DrugGym

DrugGym (www.drug-gym.org), our framework for modeling the stochastic process of drug discovery, is implemented in **Python** (version: 3.10 and above). It extends the popular **Gymnasium** library (version: 1.0.0) [102] for testing reinforcement learning (RL) algorithms.

DrugGym implements a *DrugAgent* and a *DrugEnvironment*. An experiment is run with a predefined budget, defined in terms of number of molecules made, total monetary cost, the number of DMTA cycles, or all of these. At each step, the DrugAgent receives observations and emits actions, while the DrugEnvironment does the opposite. The observations that the DrugAgent receives are the current library of molecules maintained inside the DrugEnvironment. This is an array: each row represents an individual molecule, and the columns represent all the information that is known about the molecule, such as its SMILES and other properties that may have been recorded (predictive scores and measurements). There are also columns for the current status of the molecule, which may be set to *Designed*, *Scored*, *Made*, or *Tested*, as well as the timestep when the transition was made to its current status. A molecule is Designed if it has just been ideated but no further action has been taken, Scored when the molecule is scored by every surrogate model available (with non-zero model error, called *NoisyOracles*), Made if the molecule is recorded as having been synthesized, and Tested when the molecule is run through every surrogate model available (with zero model error, called *Oracles*). Individual cells in the DrugEnvironment’s library (equivalent to the DrugAgent’s observations) may be empty, for example if a molecule has just been Designed but no additional actions have been taken.

After every step, the DrugEnvironment is responsible for ending or continuing the episode. Using its *Policy* (defined below), it assesses every molecule that has been fully Tested. If any molecule returns a utility of 1.0, the episode is ended (“terminated,” in the parlance of the Gymnasium library). An episode will also finish if the budget is exhausted (“truncated”).

### DrugAgent

DrugAgents are initialized with a Policy object, an ExploratoryStrategy object, and a sequence of action templates.

Each action template is a dictionary containing the name of the action to be undertaken (e.g., “design,” “Noisy ABL1 pIC50 docking,” “make,” “ABL1 pIC50 docking”), any parameters to associate with that assay (e.g., the number of steps to run the simulation or the ideation temperature parameter for the design step), and a batch size, which is an integer that will determine the number of molecules with which the action will be performed. The intent of this action sequence is to mimic choices that a medicinal chemist might make in a typical ADME assay cascade [53]. The DrugAgent generates an action based on the information currently available in its actions. As it does, it replaces the batch size with an appropriate number of indices, which are associated with rows in the DrugEnvironment’s library. These are the chosen compounds on which the action will be performed.

To enforce the DMTA cycle progression, the DrugAgent filters observations depending on the action that is anticipated in the next step, outlined below:

**Table S2.** Actions and Filter Criteria.

| Action Type | Filter Criteria | Reasoning |
| --- | --- | --- |
| Design | Scored OR Tested | Do not ideate analogs of molecules for which no information is known |
| Make | Scored OR Designed | Do not order a synthesis of a molecule that has already been synthesized |
| Type starting with “Noisy” (i.e., Score step) | Designed only | Do not score molecules that have already been scored, made, and tested |
| All other actions (i.e., Test step) | Made only | Molecules that have not been synthesized cannot be tested |

The DrugAgent selects these molecules using its Policy and ExploratoryStrategy. The Policy computes a “utility” for each molecule using all available information (scores, given by NoisyOracles; and measurements, given by Oracles). Briefly, our standard Policy employs a hybrid sort that includes non-dominated sorting [71] into Pareto fronts and ranking by weighted average within fronts. The Policy is described in more detail below. Given a ranking of these utilities, the ExploratoryStrategy returns a final selection. The purpose of the ExploratoryStrategy is to balance exploiting molecules that appear to be maximally valuable in the present step with the option to explore unknown but potentially promising chemical space. While our experiments do not include a step for updating our models, nor do we use probabilistic predictions as in Bayesian optimization, all of our scores and tests are associated with irreducible error, so striking a balance is likely to improve search efficiency. Here, we use an 𝜖-greedy strategy, which is a simple and interpretable option for parameterizing the tradeoff between exploitation and exploration [73].

### DrugEnvironment

DrugEnvironments are initialized with an integer budget, Policy object, list of assays (Oracles) and surrogate predictive models (NoisyOracles), and a Designer, responsible for taking molecules and returning molecules nearby in chemical space. We describe the Designer in-depth below. The DrugEnvironment parses actions from the DrugAgent. It ensures that the DMTA progression is respected by asserting conformity to the action filter criteria outlined above. As it performs the action, it annotates the value for the corresponding row in its library and also annotates a timestamp of the action. We use this timestamp for later accounting of program lengths and costs.

The choice of Policy, which is how the environment assigns utility to the molecules in its library, is slightly different from that of the DrugAgent. The DrugAgent uses a hybrid sort that incorporates non-dominated sorting, because it must rank molecules by considering tradeoffs of several competing objectives. However, the DrugEnvironment is not ranking but rather reporting the maximum utility observed. Therefore, the Policy of the environment returns a simple weighted average from the utility of each objective (this is identical to the second step of the hybrid sort).

After every step, the DrugEnvironment returns observations (its library), reward (the maximum utility observed for any individual molecule in its library for which all measurements are available), termination state (met objectives), and truncation state (exhausted budget). If either termination or truncation states are set to 1, the episode ends. This construction implements the Gymnasium API.

### Designer

Our Designer is intended to mimic standard routines of medicinal chemistry. Given a starting molecule (identified by a SMILES) and its synthesis route, we run one of two routines, inspired by previously reported ideation strategies for hit-to-lead candidates [115].

*Replace.* Given a fixed route (provided by the original molecule), we replace one or more reactants. The “new” reactant that replaces the old comes from the **Enamine** Global Stock (version June 30, 2023) [54], and the search criteria for the new reactant is the sum of Tanimoto similarity based on ECFP4/1 1024 bit fingerprints computed by **OpenBabel** (version: 3.1.1) and a size similarity score based on the number of heavy atoms in the molecules, computed by **RDKit** (version: 2024.03.1). We consider size explicitly during this search because Tanimoto similarity alone does not take it into account. We found that it will rank a building block with several identical repeats of patterns from the original reactant (e.g., a heterocycle) over one that is off by a single atom. We believe this implementation is closer to the meaning of similarity. The ranking over all of the building blocks is given by **chemfp** (version: 4.1) [116].

We re-weight the ranking with a Boltzmann distribution parameterized by an ideation temperature *T* :

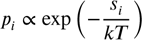

where 𝑝_𝑖_ is the new probability assignment for the 𝑖th building block, 𝑠_𝑖_ is the similarity score compared to the original molecule for the same building block, and 𝑘𝑇 is the product of the Boltzmann constant with the ideation temperature. To obtain the new ordering, building blocks are sampled without replacement from a multinomial distribution (**PyTorch** version: 2.3.0) using the values of 𝑝_𝑖_ as weights. Higher temperatures are more likely to select building blocks that are further down the ranking; lower temperatures conform to the original order. Hence, high temperatures are more adventurous. An additional parameter is the number of reactants that may be replaced as part of any given sampled molecule, compared to the original synthesis route. Since throughput is bottlenecked by reading the building blocks from an SDF on disk, we perform the replacements using lazy generators that minimize disk I/O. For multi-step reactions, we generate reactants at reaction-time, working from the “leaves” of the reaction tree toward the product. Reactions are run forward (using **RDKit** version: 2024.03.1) until sufficient products are produced. Upon failure due to reactant incompatibility, the loop continues to the next set of reactants until a sufficient number of products are enumerated.

#### Grow

The original molecule is used as one of the reactants, with the other being a randomly chosen building block. To avoid oversized products, we limit the size of the chosen building block to 10 heavy atoms. Since the appropriate reaction is unknown ahead of time, every reaction in the user-defined reaction repertoire is attempted in every loop. If none are compatible, the loop continues with the next randomly-chosen building blocks. The Grow routine in particular is necessary to approach the size needed for achieving drug-likeness [117].

All reaction products are kekulized in **RDKit** (version: 2024.03.1) and each resulting SMILES is recorded in canonical form. Other general features of the Designer include its cache, maintained so that it avoids generating the same molecule again, and an optional flag that protects non-reactive atoms from participating in subsequent reactions. This feature, and another that enforces a strict substructure match in building blocks, can be used to restrict the chemical space that can be enumerated. By default, all permutations of reactants are attempted with a reaction before moving on to the next set. Our reactions are based on SMIRKS patterns from **SmilesClickChem** (version: 1.0.1). We found that the order of the reactions matters, since some reactions tend to be more generally compatible than others. To estimate the general compatibility, we tested each reaction for compatibility against 10,000 samples of Enamine building block pairs. We reordered the reactions based on these estimates.

We are not the first to propose a synthetic enumeration system like this [118], but we have made it very efficient (**Supplementary Figure 18**).

### Oracle

Oracle functions in DrugGym are intended to be realistic surrogates of real drug discovery assays. They receive molecules and return a value for each molecule. We also implement a NoisyOracle to represent predictive models, and we call these predictive models. These wrap Oracles with an additive Gaussian error, which can be parameterized with location and spread parameters. Here, we only modify the spread, so our error distributions are unbiased relative to the original Oracle values. We now describe the different Oracle functions. ^1^

*ABL1 pIC50.* We use docking to mimic realistic structure-activity relationships that possess activity cliffs, which frequently frustrated real drug discovery programs [45]. For ligand preparation, we employ **RDKit** (default settings, version: 2024.03.2) for protonation and conformer generation, and **Meeko** (version: 0.5.0) for computing charges (default settings except `rigid_macrocycles=True`). We do not perform target preparation ourselves, instead using a prepped ABL1 pocket fragment given in **DOCKSTRING** (version: 0.3.2) [119]. We adopt their pocket center, which is `(15.851, 14.647, 3.904)`. For docking, we make use of **uni-dock** (version: 1.1.2 with conda installation), a GPU-accelerated implementation of **Autodock Vina** [56, 120]. We use the “detailed” setting, which sets exhaustiveness to `512` and max-step to `40`. All other settings use defaults. **Uni-dock** returns a docking score in terms of Δ𝐺, but we wish to interpret results in terms of pIC50. Therefore, we implemented a conversion to pIC50, which involves multiplying the docking score (which is in terms of Δ𝐺 kcal/mol) by a coefficient.

To derive that coefficient, we note that 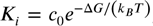, where *c*_0_ is a reference concentration (here, 1 M); we assume the enzyme obeys Michaelis-Menten kinetics, allowing us to relate the half maximal inhibitory concentration (IC50) to the inhibition constant *K_i_* using the Cheng-Prusoff equation:

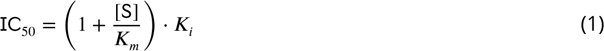

We assume that our assay has [𝖲] << 𝐾_𝑚_, such that IC50 ≈ 𝐾_𝑖_). The definition of pIC50 is − log_10_(𝖨𝖢50∕𝑐_0_). Then,

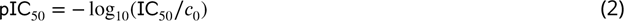

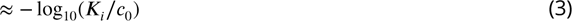

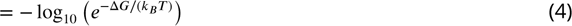

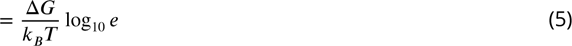

At room temperature ( *T* = 300 Kelvin), *k_B_T* ≈ 0.596 kcal/mol.

*Log S.* We trained a **CatBoost** model (version: 1.2.5) [121] on **AqSolDB** [61]. Like ChemProp-RDKit [122], we generate input features for regression using RDKit, run with multiprocessing via **scikit-mol** (version: v0.2.0) [123]. These features are: ExactMolWt,FpDensityMorgan1,FpDensityMorgan2,FpDensity Morgan3,HeavyAtomMolWt,MaxAbsPartialCharge,MaxAbsPartialCharge,MinAbsPartialCharg e,MinPartialCharge,MolWt,NumRadicalElectrons,NumValenceElectrons,MolLogP,Fraction CSP3,HeavyAtomCount,NHOHCount,NOCount,NumAliphaticCarbocycles,NumAliphaticHeteroc ycles,NumAliphaticRings,NumAromaticCarbocycles,NumAromaticHeterocycles,NumAromati cRings,NumHAcceptors,NumHDonors,NumHeteroatoms,NumRotatableBonds,NumSaturatedCarb ocycles,NumSaturatedHeterocycles,NumSaturatedRings,RingCount. Using a uniform sampling strategy, we split these data into 80% train and 20% test, and our model recorded 0.784 MAE on the test set (**Supplementary Figure 13**). Our results compare favorably to the current leaderboard on the Therapeutics Data Commons [36].

*Log P.* We use **RDKit**’s MolLogP descriptor, which is an implementation of Crippen’s method [58].

### Utility function

The utility function maps from the values received from an Oracle to (−∞, 1], expressing the relative benefit that can be gained from selecting a molecule with that value. This mapping is determined by a user-defined target candidate profile (TCP), which is a set of threshold pairs (an upper and lower value to signify acceptable and ideal ranges). For all experiments, we use a challenging but not overly difficult TCP that includes ABL1 pIC50, log S, and log P (**Supplementary Table 3**, reproduced from the **Introduction**).

**Table S3.** Target candidate profile (TCP) objectives used in this work. Associated methods used here for computing corresponding Oracles in DrugGym are noted.

| Objective | Units | Oracle Method | Dynamic Range | Thresholds |  |
| --- | --- | --- | --- | --- | --- |
|  |  |  |  | Acceptable | Ideal |
| Affinity | pIC50 | Docking [56] | [3, 11] [57] | [8, ∞) | [9, ∞) |
| Lipophilicity | Log P | RDKit [58] | [-1, 8] [59] | [0, 4] | [0, 3] |
| Solubility | Log S | Gradient Boosted Tree [60] | [-8, 1] [61] | [-4, 0] | [-3, 0] |

The corresponding oracle classes are DockingOracle, CatBoostOracle, and RDKitOracle, described above. Given the TCP, we define a piecewise multi-dimensional utility function as follows:

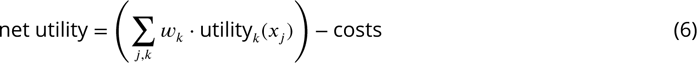

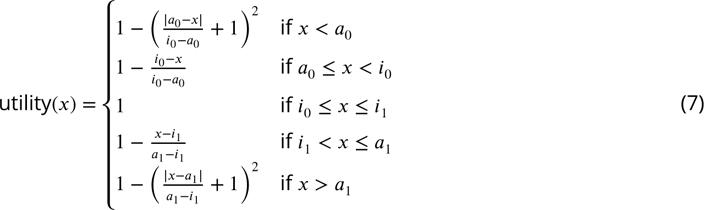

This functional form was chosen because it implies that any value within the ideal range is equally valuable, and that no molecule can be more valued than those within the ideal range; that there is a monotonic increase in value for molecules that fall between the acceptable and ideal ranges; and that values whose properties fall outside of the acceptable ranges are proportionally penalized. In this case, with a quadratic penalty common to constrained optimization problems [68]. Our goal is to find a molecule that maximizes utility.

### Policy

The Policy is designed to return a scalar that can be used by the DrugAgent to select molecules. We draw a distinction between values returned by Oracles (“measurements”) and those returned by NoisyOracles (“scores”). By construction of the action filtering criteria discussed in DrugAgent, we will have scores for any molecule that has measurements, but not the converse. Therefore, we implemented a graceful fallback, which uses measurements when they are available, but otherwise uses scores in their place. To implement a well-defined metric space, molecules can only be compared if the resulting values are complete, by which we mean that every “assay” has an associated score or measurement.

To compose utilities across multiple objectives, we use one of two methods. The first and simpler is weighted average (alternatives here include min, max, sum, and product, compared in **Supplementary Figure 18**). The weights are user-defined. Here, we use 0.8 for pIC50, 0.1 for log S, and 0.1 for log P, and we do not deviate from these in any of our experiments. It is not necessary that they add up to 1. We chose these weights to influence the optimization, but different weights could easily be chosen, or even optimized; a weights schedule that changes according to traditional assay tiers is also possible. The weighted average sort is used to obtain the current reward in the DrugEnvironment. We also implement a hybrid sort, which is used in the DrugAgent. Our hybrid sort uses NSGA-II (**pymoo** version: 0.6.1) to define non-dominated fronts. We use the same weighted average as before to break the tie within fronts. These rankings are composed using `numpy.lexsort` (**NumPy** version: 1.26.4). We chose this criteria because it satisfies Pareto optimality [69] and minimizes deviation from the ideal range of the TCP.

After several rounds of DMTA, we will have some molecules with complete measurements, and others that have only been scored. Which molecules should serve as the basis for designing analogs in subsequent DMTA cycles? One rule is to ideate around tested molecules only. However, this severely limits the search space and may get trapped in local minima, especially in the early part of campaigns. We sought a procedure for selecting from scored and tested molecules on equal footing. While we expect the Gaussian error of our predictions to be homoscedastic, this is not true under selection. In that case, the ranking of observations considers both the underlying “true” values and the noise. Hence, the distribution of this subset will systematically overestimate the true value with high probability (**Supplementary Figure 17**). To address this, we employ a simple linear correction that uses available measurements and corresponding predictive scores from molecules that have been tested. At every time step, we fit an OLS regressor (**statsmodels** version: 0.14.1) that maps from predictions to actual measurements. This doesn’t improve predictive power, but it does rescale scores so that predictions and actual measurements can be directly compared in the selection step. We employ the OLS correction only after 30 molecules have been tested, so sample statistics have approximately converged. The correction is applied only to scored molecules (not actual measurements), and these corrected scores are used in the utility function transformation.

### ExploratoryStrategy

Given the utility predicted by the Policy, the DrugAgent must choose the indices of molecules to progress into the next DMTA cycle action. We call this function the ExploratoryStrategy. We use the classic 𝜖-greedy method, because it is a competitive method that trades off exploration and exploitation in an interpretable fashion [73]. Another simple alternative would be Boltzmann exploration [124], but other ExploratoryStrategies could be adopted as part of a more sophisticated DrugAgent architecture employing neural networks and search algorithms.

### Experimental Defaults

*Hit selection.* The standard setting of the experiments begins when 5 starting hits are selected from the SDF of **Enamine**’s DSI-poised Library (version: 2 March 2021) [125]. This selection is restricted to the library subset that is compatible with the two-reactant, one-step reaction repertoire from **SMILESClickChem** (version: 1.0.1) [55]. Reaction compatibility is determined by non-zero product yields from permuting the reactant ordering with forward synthesis in **RDKit** (version: 2024.03.1) [126].

*Action sequence.* A sequence of action templates determines what action will be generated next by the DrugAgent.

**Table S4.** Baseline action sequence for all experiments. Every product generated in the Design step is scored in the Score step that immediately follows.

| Sequence | Action Type | Batch Size | Parameters |
| --- | --- | --- | --- |
| 1 | Design (Replace) | 8 | Products: 5, Temperature: 0.16 |
| 2 | Score | 40 | Noisy ABL1 pIC50, Noisy Log P, Noisy Log S |
| 3 | Design (Grow) | 8 | Products: 5 |
| 4 | Score | 40 | Noisy ABL1 pIC50, Noisy Log P, Noisy Log S |
| 5 | Make | 8 |  |
| 6 | Test | 8 | ABL1 pIC50, Log P, Log S |

Every product generated in the Design step is scored in the Score step that immediately follows it. The effect of this default is that 10 compounds are scored for every one that is made and tested. Our 𝜖-greedy uses 𝜖 = 0.2, meaning that 20% of the time, a compound was chosen at random (without replacement) rather than according to the utility provided by the Policy.

### Chemical properties

We ran 100 trials using the default parameters outlined above (with 𝜎 = 1.0 used in the NoisyOracles surrogate models of each of our TCP objectives). Due to measurement error, stochasticity in decision-making, and the varying quality of starting hits, programs complete at different times. To make an appropriate comparison across compounds at the beginning, middle, and end of campaigns, we create a normalized measure of campaign progress. Following Beckers et al. [79], we measure progress as the percentage of compounds that have been “made” in the simulation. We use progress percentage to study longitudinal impacts of different ideation and selection strategies.

To generate **Figure 4B**, we first compute cumulative progress percentage assignments. That is, for each progress percentile, we gather all the molecules that have had this progress percentage or lower (i.e., 100% cumulative progress should be very smooth, since it contains *all* molecules in the series). We pool this analysis across every trial. Since we are interested in the right tail of the distribution, we compute the cumulative distribution (CDF) for molecules that can be found at a given ABL1 pIC50 value or *greater*, stratified by the cumulative progress percentage just described.

Figure 4C plots the *average* of various computed properties on the same dataset across the progress percentage (not cumulative), with 99% bootstrapped confidence intervals. These properties include: ABL1 pIC50, Log P, Log S, HeavyAtomCount, RingCount, FractionCSP3, LiPE, LE, and QED. The first three were computed using our DrugGym Oracles as described before. The rest were computed using **RDKit** (version: 2024.03.1).

Figure 4C visualizes the vertices and edges of the design lineage graph for molecules that were made within a single representative trial. We constructed the graph using **networkx** (version: 3.3) [127] and used the Reingold-Tilford algorithm implemented in **python-igraph** (version 0.11.5) [128] to generate coordinates. Edges are cubic Bezier curves. The color intensity is given by the utility function for the corresponding TCP measures from the compounds.

### Chemical structures

In Figure 5A, the same normalization procedure as described in the previous section was used to compute progress percentage. We used the same 100 trials with 𝜎 = 1.0 for the NoisyOracles. We pooled all the Made molecules and segmented them by progress percentage. We then sampled molecules randomly and visualized them using **RDKit** (version: 2024.03.1).

The synthetic route in Figure 5B was created by selecting one of the best-performing molecules across all of our simulations and using the `dgym.molecule.Molecule.dump` method to obtain its synthetic route (referring to “dumping” or serializing a data model). The synthetic route is preserved from when the molecule was generated by the Designer. It includes SMILES, building block identifiers from **Enamine**, and reactions. An example synthetic route dump:

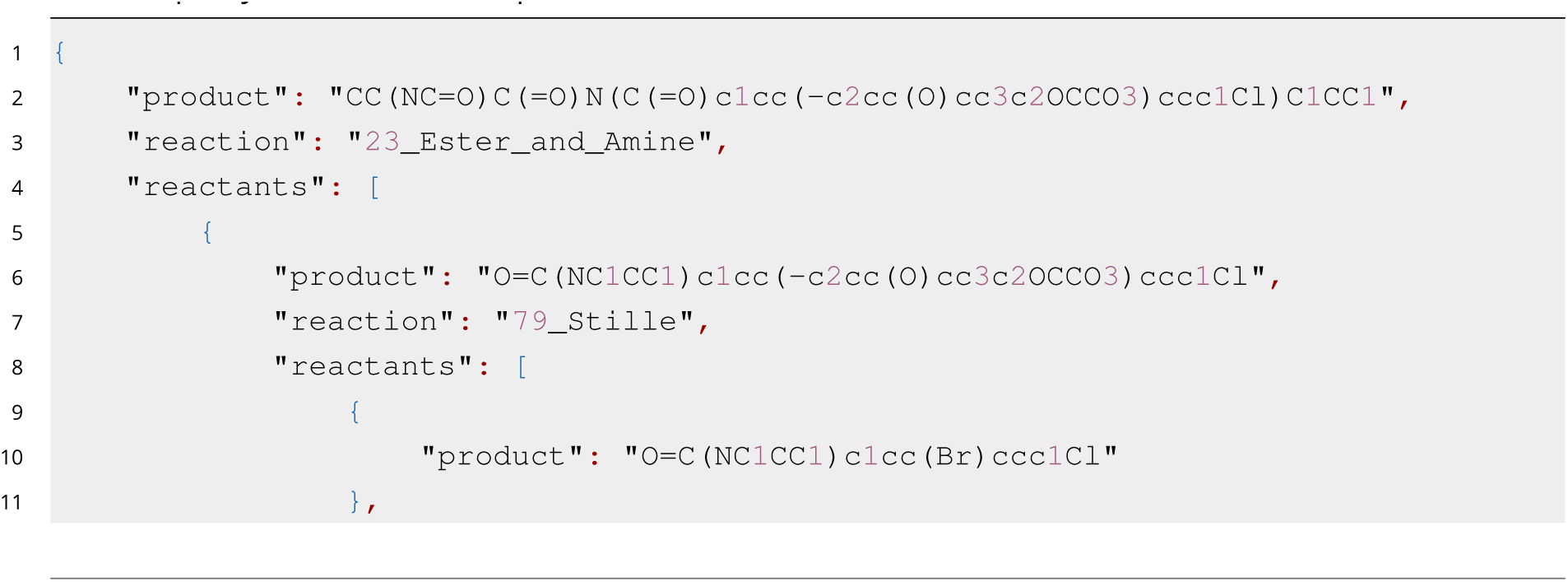
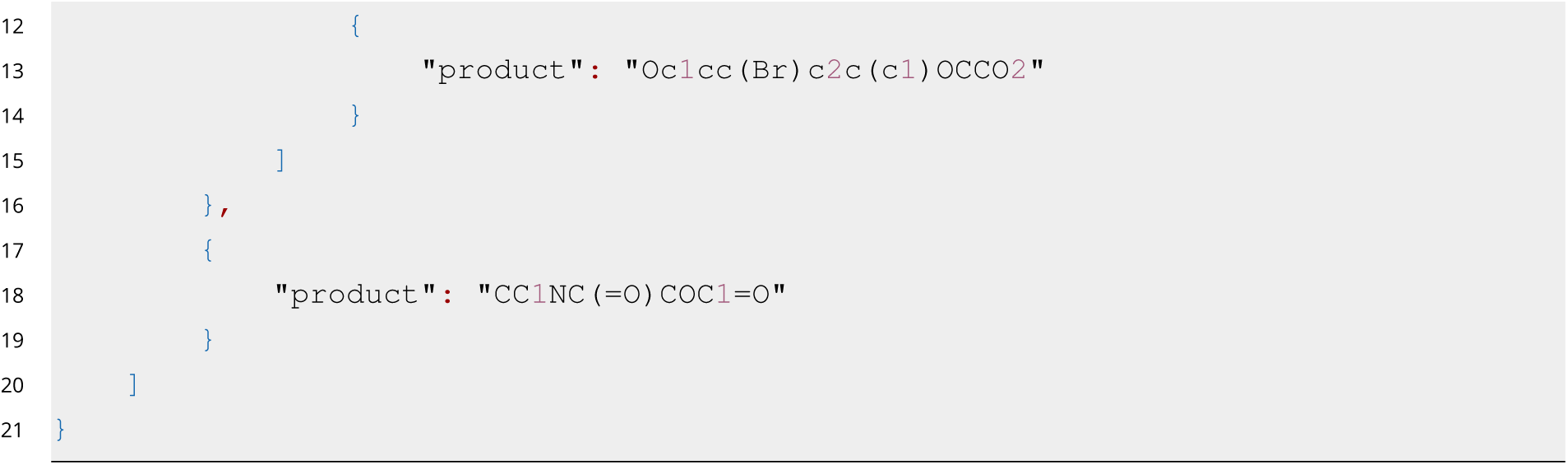

We visualized these molecules in **RDKit** (version: 2024.03.1).

The molecules in Figures 6 and **7** were also visualized in RDKit, and their properties were computed using DrugGym Oracles as described previously. The DMTA cycle value is that step assigned to the molecule when it was synthesized (assigned Made status).

### Ideation creativity

Figure 8A is a conceptual mockup based on a blog from Greg Landrum [129], intended to illustrate the Designer routines described earlier.

Figure 8B was created by repeated use of the DrugGym Designer: first, two **Enamine** building blocks were chosen at random. Then, all the reactions from **SmilesClickChem** were tested for compatibility with these reactants. The first to prove compatible was used to construct a product. This process was repeated with one more building block (using the Designer’s Grow routine). Finally, we apply the Designer’s Replace mode with different setting of ideation temperature `0.00, 0.04, 0.16`, and `1.00` across 1, 2, and 3 reactant replacements. The resulting products were visualized using **RDKit** (version: 2024.03.1).

For Figure 8C, we use a batch size of 24 rather than the usual batch size of 8. However, this batch size was used across all the trials and experimental settings shown in this figure. We use **lifelines** (version: 0.27.8) to compute a Kaplan–Meier estimator of the probability that an experiment had not yet terminated successfully by a given cost, such as the number of molecules made. We use this estimator because it was designed to be robust to right-censorship, as can occur with very long-running simulations like ours. We compute the Kaplan-Meier estimator for trials performed with escalating ideation temperatures: `0.0`, `0.02`, `0.04`, `0.08`, `0.16`, `0.32`, and `0.64`. In addition, each of these ideation temperature settings was tested with a different maximum of reactant replacements: 1, 2, or all reactants in the synthetic route. To derive the plot shown, we find the Kaplan-Meier estimate of success rate corresponding to a 3200 molecule budget. These success rates represent the proportion of trials that are expected to succeed within 3200 molecules *or fewer*. The 68% confidence interval calculated by Greenwood’s Exponential formula is represented by the shaded region.

The same analysis is extended in Figure 8D, except that the targeted quantity is 0.95 success rate. We use the Kaplan-Meier estimator at or above 0.95 success rate to find costs. These should be interpreted as the “prices” necessary to provide guarantees for given probabilities of success. They are not estimates of the *average* cost.

### Model error

In this experiment, we held all variables constant except for the 𝜎 associated with the NoisyOracles (ideation temperature was set to 0.16). We ramped 𝜎 from `0.0` to `2.0` in increments of `0.5`. As before, all experimental settings were conducted with 100 replicates.

**Figures 10C and D** are identical to the two ideation creativity experiments described just above, except that the exogenous variable is model error (*σ*).

However, **Figure 10C** contains an additional setting. Here, we replaced the usual NoisyOracles entirely with GaussianOracles. These return samples from a Gaussian distribution regardless of the input. These GaussianOracles are initialized with a very large spread (𝜎 = 10^6^) centered on the mean value of each objective from past experiments. In this setting, sigma far outstrips the dynamic range of the data. We created this experiment to stress the performance degradation from overwhelming noise (or simply uninformative models). This does not mean that we stopped scoring. Rather, this experiment maintains the ratio of 10 molecules scored for every 1 tested, but those scores are not predictive of actual measurements whatsoever.

Figure 10B simply visualizes the inverted survival curves and confidence intervals from **lifelines** (version: 0.27.8), where the duration is the number of molecules that have been made and tested in the campaign.

For the exponential regression in **Figure 10E**, we use the **lifelines** Kaplan-Meier estimator to find costs associated with 2000 linear increments spanning 0.00 to 0.80 success rate. We use this estimator because it takes right-censoring into account. The costs should again be interpreted as the minimum spend necessary to achieve that success rate. By inspection of the cost curves, we proposed a 4-parameter exponential fit:

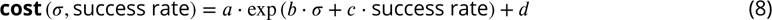

We fit this equation to the costs using `scipy.optimize.curve_fit` in **SciPy** (version: 1.13.0). The fitted parameters were `(16.65396258, 1.07881681, 2.13219256, 18.28661156)`. We again used **SciPy** to find the Pearson correlation coefficient, and we used **NumPy** to find the mean absolute error.

The last panel, **Figure 10F**, asks what would be the marginal benefit of each calculation made by predictive models, in terms of savings from finishing programs sooner. To do so, we compared our model error experiments to the case of no scoring whatsoever. We reason that this is equivalent to the medicinal chemist selecting compounds for synthesis based on their expertise alone. Formalizing this intuition, we found the median dollar cost from 100 replicates with the same experimental settings except that every compound that is ideated is made and tested (scoring models are not used in selections). Using a reported average price of $3000 for compounds that are custom-made by CROs[87], we found the median dollar cost of these trials, which was $1,812,000.00 (604 molecules made).

Across 10,000 bootstrap samples, we found the means and 95% confidence intervals of the difference between that median cost and the corresponding dollar costs for every trial (i.e., the savings), divided by the number of molecules that were scored during the trial. These represent the marginal value gained per calculation. We shaded the region of the chart where the experimental error (the expected replicability of the underlying Oracles sans Gaussian noise). The irreducible experimental error may explain why the marginal value per calculation does not rise as dramatically at the minimal error rate.

### Batch size

Here, batch size refers to the number of molecules that participate in an action in each step of the DMTA cycle. The exception is the Score step, which we hard-coded to be 5 times the batch size, chosen to correspond to the number of analogs that were ideated during Design steps. In other words, every compound that is ideated is scored. Since there are two rounds of ideation and scoring in each DMTA cycle, the net effect is that the number of molecules designed and scored is 10 times that of molecules made (80:8). We call this a 10:1 *scoring ratio*. To illustrate the effect of batch size on these numbers: if the batch size were 96 instead of 8, the corresponding number of molecules scored within each Score step would be 96 ⋅ 5 = 480. For these experiments, we use batch sizes based on the size of rows and columns in a typical microwell plate, `(8, 16, 24, 48, 96, 192`. For the same reason, these are typical batch sizes of synthesis orders and experiments.

**Figures 9A and B** represent the same analysis as that of **Figure 10B**, except that the intervention is the batch size of each trial and the model error is fixed at 𝜎 = 1.0. The other difference is in Figure 9B, for which the cost of each trial is measured in the number of DMTA cycles (indexing the timestep at which molecules were made), rather than the number of molecules made.

Figure 9C uses the same analysis of expected costs visualized in **Figure 10D**, which finds the Kaplan-Meier estimator costs that correspond to a given success rate. Here, we assign costs, in terms of both the number of molecules made and the DMTA cycles (as drawn in the previous two panels). The final visual stratifies these multidimensional costs across success rates.

### Scoring ratio

As described in the previous section, the scoring ratio refers to the relative proportions of molecules scored to those that were tested. Using the standard experimental design with batch size 8, we varied the scoring ratio at intervals between 1 and 20. A scoring ratio of 1 indicates that scores were irrelevant to selection, since every molecule that was scored was tested. The analysis in Figure 11 was followed as previously described.

### Statistics

Cumulative distribution functions are computed with inverted survival regression, with a trial’s duration given by the number of molecules that were made or the timestep of the last molecule made. We use **lifelines** (version: 0.27.8) to compute this curve and 68% confidence intervals (via Greenwood’s Exponential formula).

### Visualizations

Unless noted otherwise, all plots were created using **seaborn** (version: 0.13.2).

### Code availability

The Python code used to produce the results discussed in this paper is distributed open source under MIT license https://github.com/choderalab/drug-gym.

### A. Supplementary Figures

**Figure S12.**
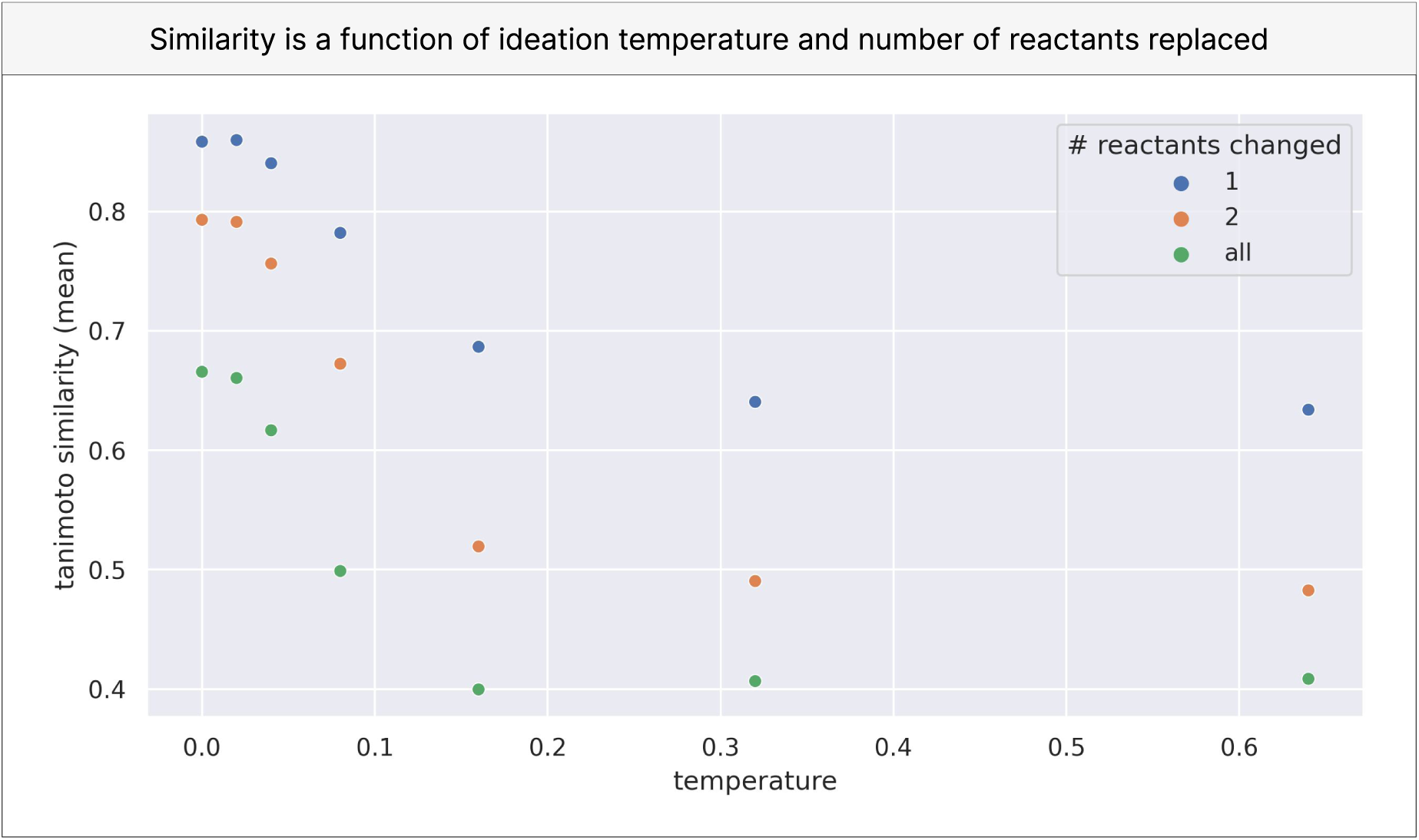
Tanimoto similarity is a function of ideation temperature and number of reactants replaced. Mean Tanimoto similarity of 500 products (each yielded from a different 2-step reaction) compared to analogs generated with different settings of ideation temperature and numbers of reactants replaced.

**Figure S13.**
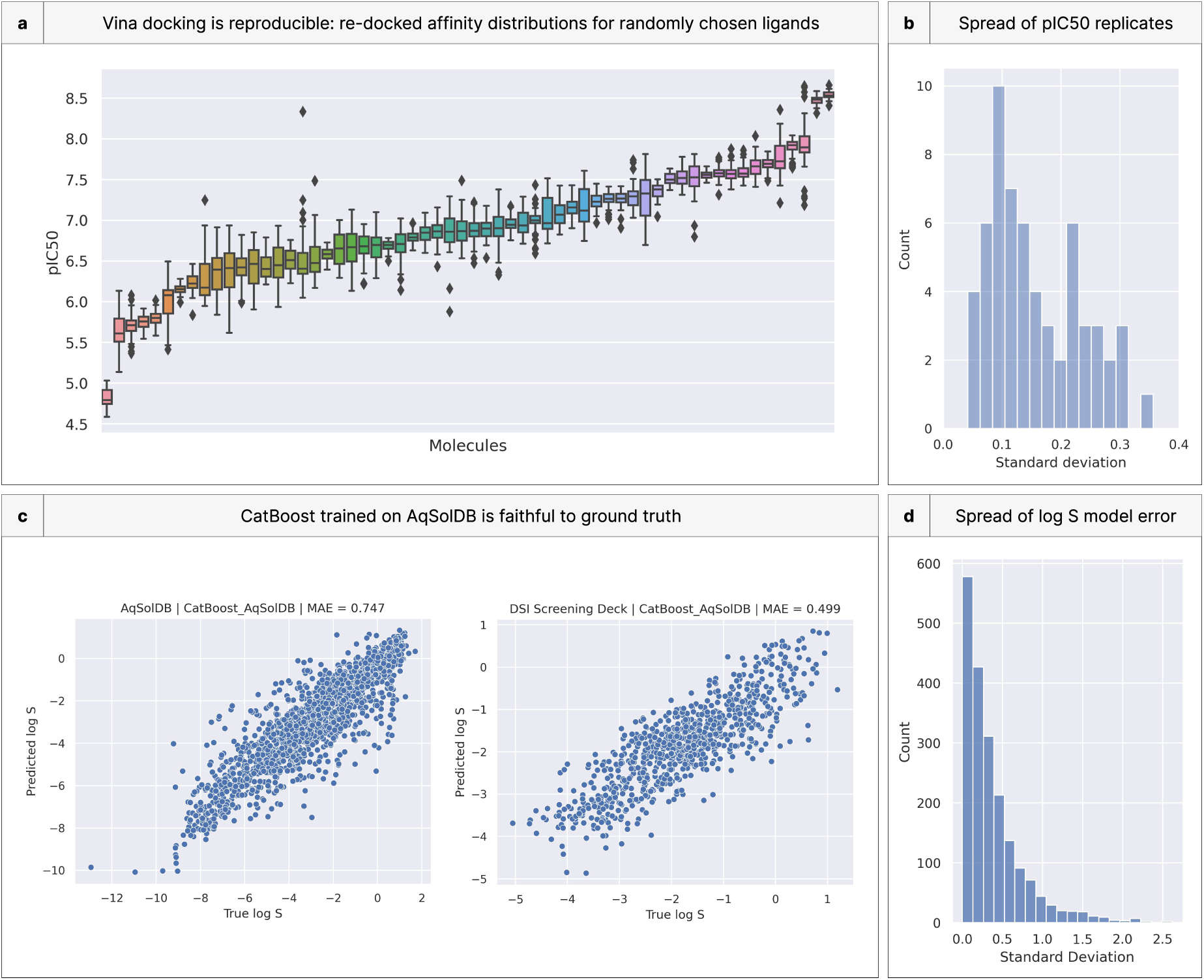
Scoring models have small but non-zero irreproducibility. **(a)** Distribution of pIC50 from re-docking 60 ligands (each generated from randomly selected Enamine building blocks). **(b)** The distribution of standard deviations from these ligands. **(c)** Scatterplots of a CatBoost regressor trained on AqSolDB [61], a well-curated solubility dataset, when compared to the held-out test set of AqSolDB and the log S reported for the DSI screening deck. **(d)** Spread of CatBoost model error compared with ground truth.

**Figure S14.**
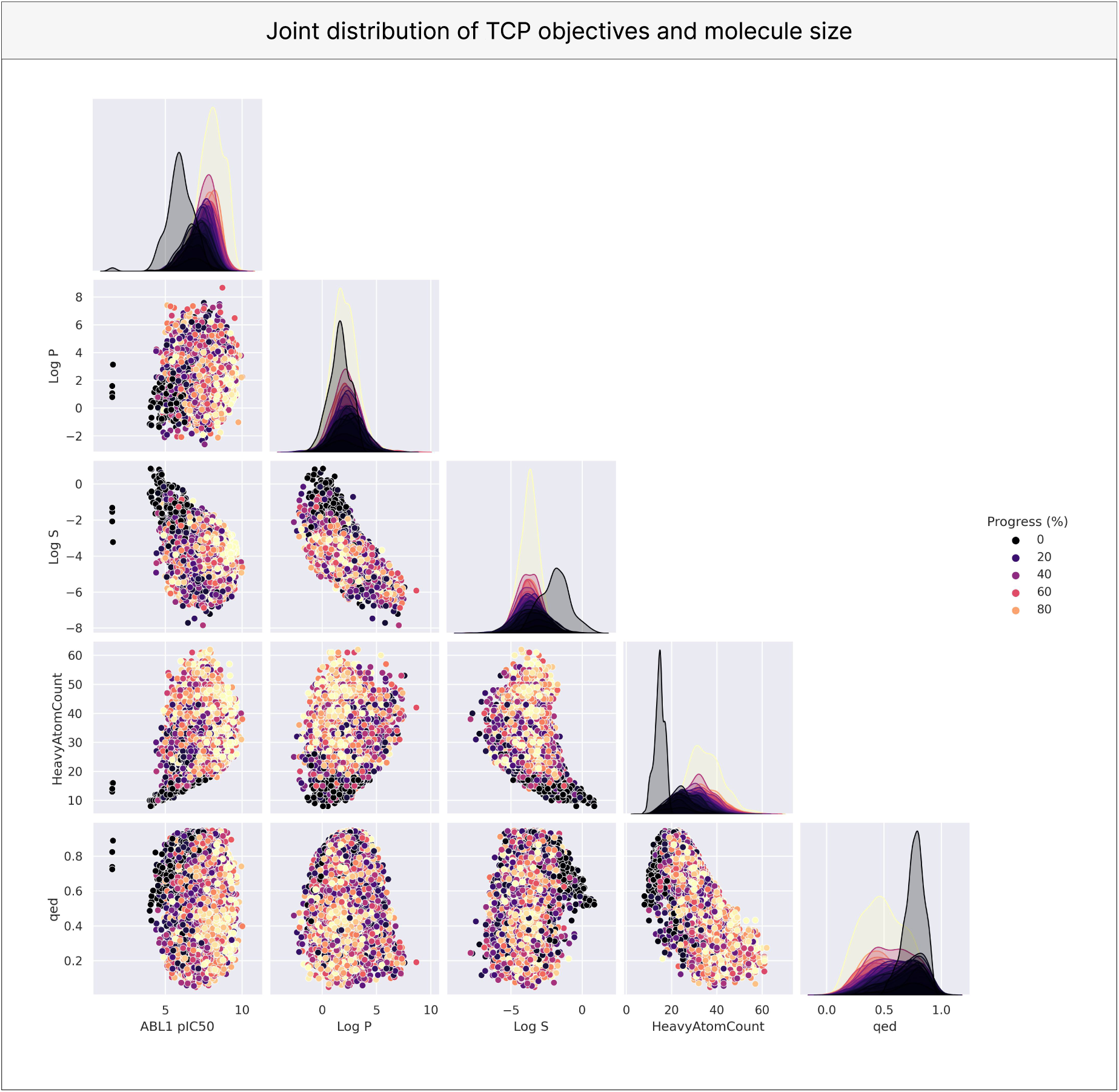
Joint distributions of various chemical properties across 100 simulated drug discovery campaigns.

**Figure S15.**
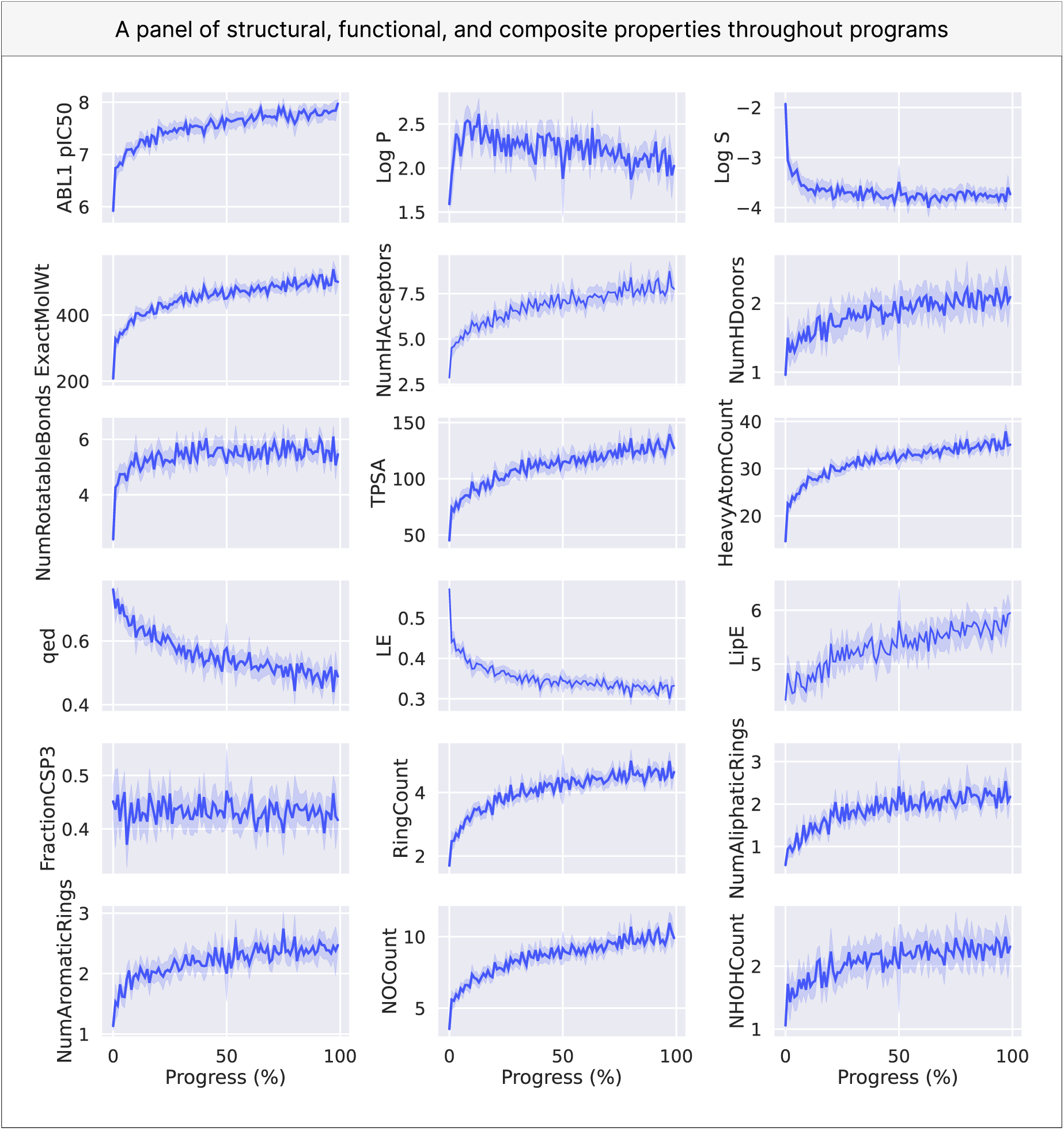
Scoring models are most useful earlier in discovery programs. Time-dependent averages of chemical properties indicative of drug-likeness averaged or many realizations of the discovery program. The first row represents the TCP objectives. With the exception of LE and LipE, the rest are measured in RDKit (acronyms after the first row, from left to right: NumHAcceptors: number of atoms in the molecule that are hydrogen-acceptors; NumHDonors: number of atoms in the molecule that are hydrogen-donors; TPSA: topological polar surface area; FractionCSP3: the ratio of sp hybridized carbons over total carbons; QED: quantitative estimation of drug-likeness; LE: ligand efficiency, pIC50 divided by the number of heavy atoms; LipE: lipophilic efficiency, difference of pIC50 and Log P; NOCount: number of Nitrogens and Oxygens; NHOHCount: number of N and O atoms with a covalent bond to a hydrogen).

**Figure S16.**
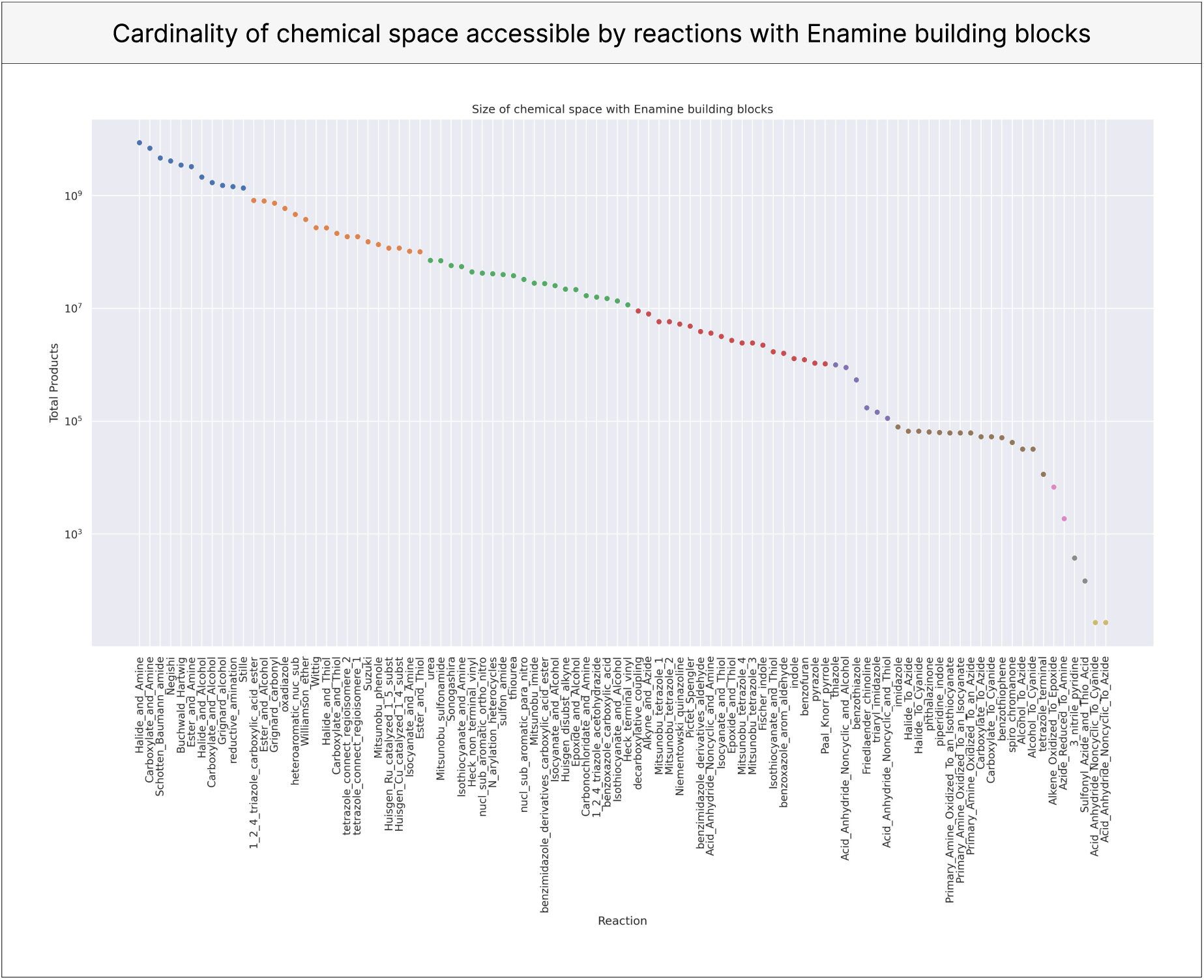
The accessible chemical space in DrugGym is vast. For each reaction in the SMILESClickChem set, the SMIRKS of its reactants was compared with every building block in the Enamine global stock for substructure match. The product of these values is the maximum possible number of products that can be synthetically enumerated using the reaction and building blocks.

**Figure S17.**
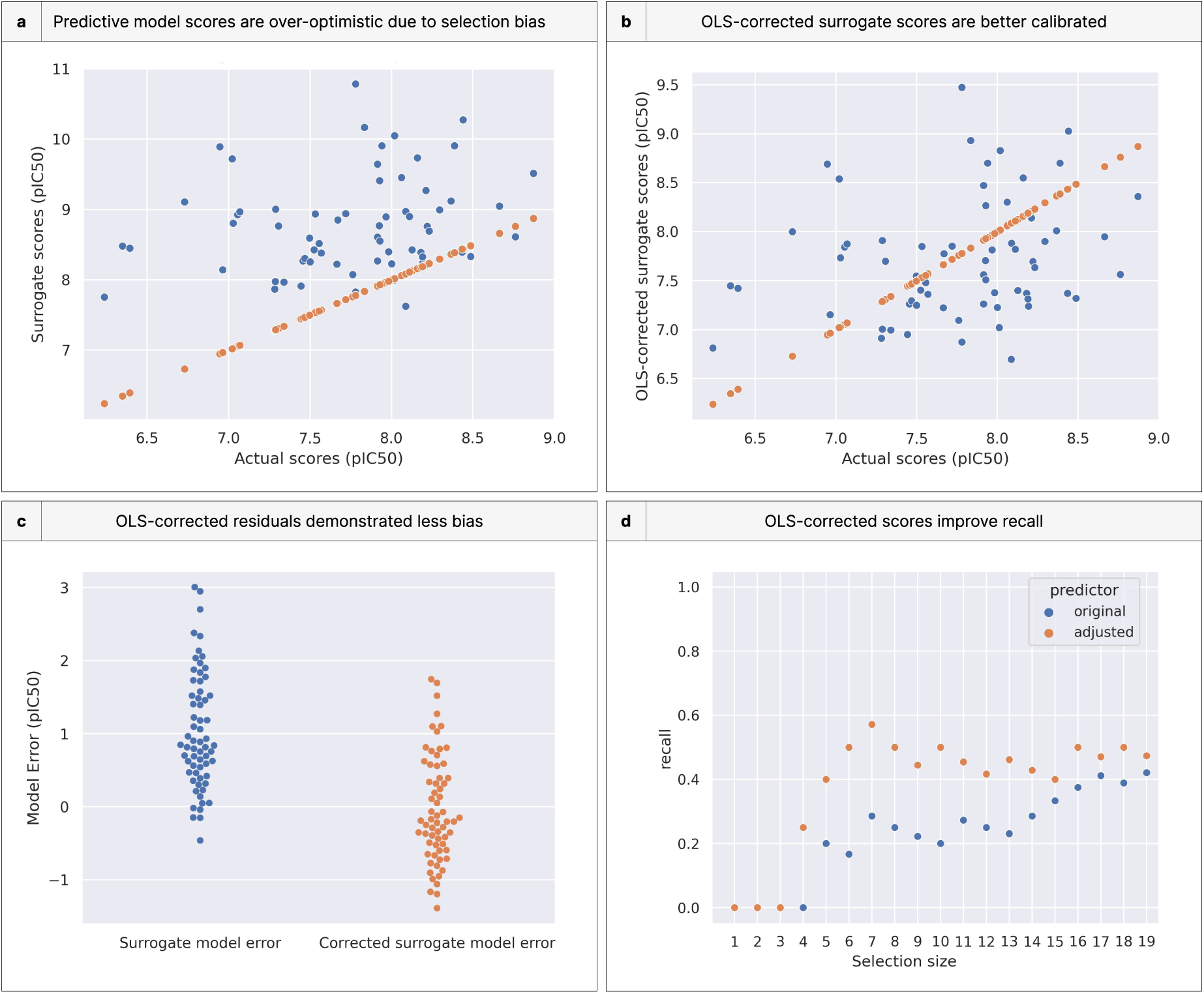
An OLS correction effectively calibrates predictive model scores. **(a)** A campaign was simulated as usual with *σ* = 1.0. ABL1 pIC50 scores from this campaign were compared to the “true” ABL1 pIC50 Oracle measurements, indicating that selecting for the right tail of noisy scores finds a set of molecules with systematically over-estimated ABL1 pIC50. Orange dots are the actual values and blue are the surrogate scores. **(b)** Using the OLS-adjusted surrogate scores results in visibly better calibration. Orange dots are the actual values and blue are the surrogate scores. **(c)** Compared with the residuals of unadjusted scores, where almost all are positive, the residuals of OLS-adjusted scores are centered on zero. **(d)** The top 𝑘 molecules as derived from unadjusted and adjusted scores are compared for recall against the “ground truth” top 𝑘 molecules. OLS-adjusted scores have superior recall.

**Figure S18.**
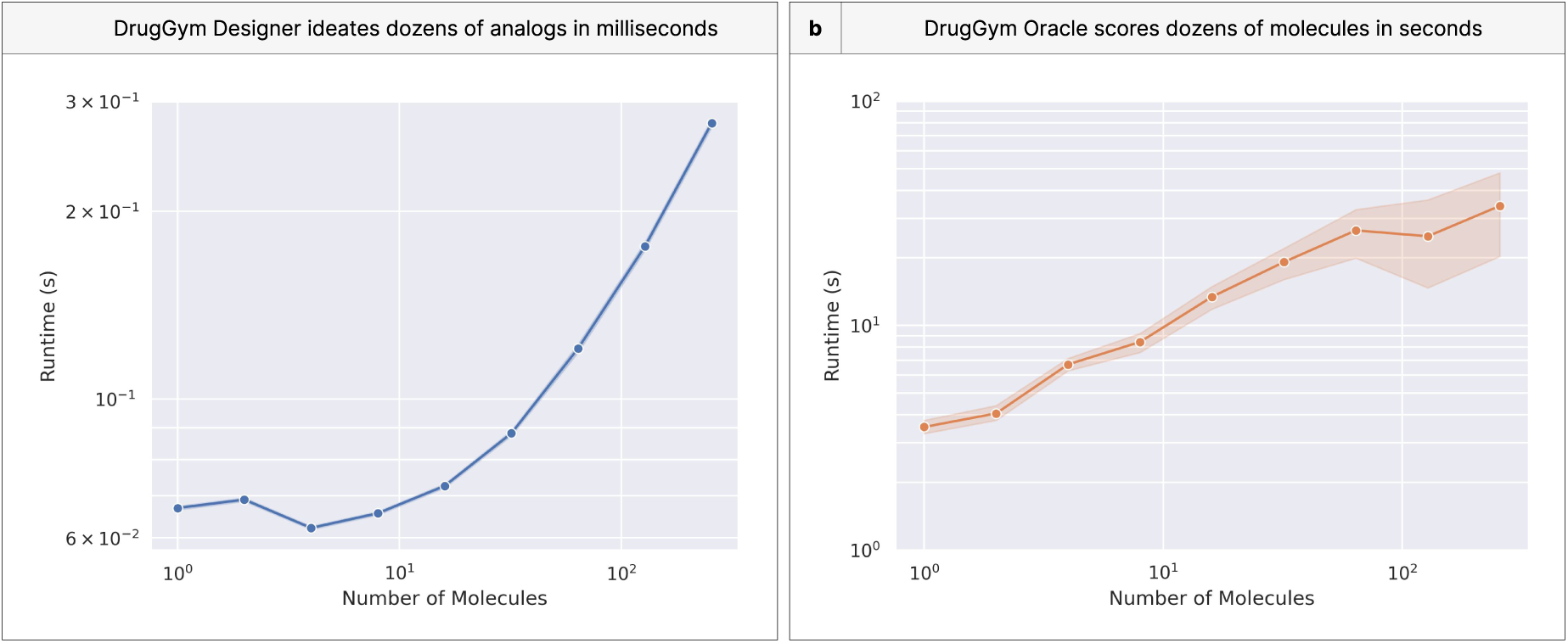
DrugGym routines run efficiently. Mean and 95% confidence interval for the runtime of various DrugGym routines. **(a)** DrugGym Designer can ideate dozens of analogs in milliseconds. Analogs were designed around a product of two randomly chosen Enamine building block reactants (𝑛 = 1000). **(b)** DrugGym Oracles can score dozens of molecules in seconds. Analogs generated in the same fashion as in the Designer runtime experiment were scored using DrugGym’s standard TCP Oracles: ABL1 pIC50 docking with uni-dock, Log S with CatBoost trained on AqSolDB, and Log P using MolLogP in RDKit (𝑛 = 100). **System specifications:** 13th Gen Intel(R) Core(TM) i9-13900H 2.60 GHz with 32GB RAM, NVIDIA GeForce RTX 4070 Laptop GPU with CUDA V11.7.64, Ubuntu 22.04.2 LTS.

## Footnotes

1 Our convention is to prefix “Noisy” to the names of NoisyOracles (that of “ABL1 pIC50” would be “Noisy ABL1 pIC50”), and we use the term “assay” to refer to the task they share with their corresponding Oracle (in this case, “ABL1 pIC50”).

## Notes

http://www.drug-gym.org

https://github.com/choderalab/drug-gym

